# Identification of Mfn2-S249 as a Phosphoregulatory Switch of Mitochondrial Fusion Dynamics

**DOI:** 10.1101/2022.01.11.475884

**Authors:** Sanjay Kumar, Aaron Ramonett, Tasmia Ahmed, Eun-A Kwak, Paola Cruz Flores, Hannah R. Ortiz, Paul R. Langlais, Karthikeyan Mythreye, Nam Y. Lee

**Affiliations:** Division of Biology, Indian Institute of Science Education and Research, Tirupati 517507, India; Department of Pharmacology, University of Arizona, Tucson, AZ 85724, USA; Department of Chemistry & Biochemistry, University of Arizona, Tucson, AZ 85724, USA; Department of Medicine, University of Arizona, Tucson, AZ 85724, USA; Department of Pathology, University of Alabama at Birmingham, 35294, USA; Comprehensive Cancer Center, University of Arizona, Tucson, AZ 85724, USA

**Keywords:** Mitochondrial dynamics, TGF-β, TAK1, Mfn2, white adipocytes, cell metabolism

## Abstract

Mitochondrial remodeling is a fundamental process underlying cellular respiration and metabolism. Here we report TAK1 as a direct regulator of mitochondrial fusion. TAK1 is activated by a variety of mitogenic factors, cytokines and environmental stimuli, which we find induces rapid fragmentation through Mfn2 inactivation. TAK1 phosphorylates Mfn2 at Ser249, which inhibits the binding of GTP required for Mfn trans-dimerization and mitochondrial membrane fusion. Accordingly, expression of Mfn2-S249 phosphomimetics (Mfn2-E/D) constitutively promote fission whereas alanine mutant (Mfn2-A) yields hyperfused mitochondria and increased bioenergetics in cells. In mice, Mfn2-E knock-in yields embryonic lethality in homozygotes whereas heterozygotes are viable but exhibit increased visceral fat accumulation despite normal body weight and cognitive/motor functions compared to wildtype and Mfn2-A mice. Mature white adipocytes isolated from mutant mice reveal cell-autonomous TAK1-related effects on mitochondrial remodeling and lipid metabolism. These results identify Mfn2-S249 as a dynamic phosphoregulatory switch of mitochondrial fusion during development and energy homeostasis.

## INTRODUCTION

Mitochondria are dynamic organelles that undergo continuous fusion and fission to regulate cellular respiration, metabolism and stress response (Eisner et al., 2018; Giacomello et al., 2020; Youle and van der Bliek, 2012). In mammalian systems, the core components comprise Drp1, a cytosolic GTPase recruited to the mitochondria to mediate fission, and mitofusin-1 and −2 (Mfn1 and Mfn2) and Opa1, two classes of transmembrane GTPases that promote the outer- and inner-membrane fusion, respectively (Chen et al., 2003; Olichon et al., 2003; Smirnova et al., 2001). Relative to fission, less is known about mechanisms governing fusion although the two structurally-related Mfns are believed to initiate the tethering of opposing mitochondrial membranes by forming either homotypic or heterotypic dimers (Cao et al., 2017; Li et al., 2019; Qi et al., 2016). Current evidence supports the binding of GTP as a crucial first step of *trans* Mfn dimerization that precedes the GTP hydrolysis-induced conformational changes necessary to bring the outer- and inner-mitochondrial membranes into closer proximity (Li *et al*., 2019).

A fundamentally elusive aspect of mitochondrial fusion relates to how posttranslational modifications (PTMs) affect the intrinsic catalytic functions of the fusion machinery. In the case of Mfn1 and Mfn2, several distinct phosphorylation, ubiquitination and acetylation sites have been identified although most, if not all, existing PTMs appear to modulate Mfn protein stability/degradation (Chen and Dorn, 2013; Dagda et al., 2009; Leboucher et al., 2012; Park et al., 2014; Pyakurel et al., 2015; Zhou et al., 2010). And while PTM-induced changes in Mfn expression are often closely linked to apoptosis or mitophagy in damaged mitochondria, they nevertheless represent largely slow-acting and bioenergetically costly mechanisms of mitochondrial remodeling. Hence, it is unclear whether PTMs can exert more precise, dynamic and reversible changes in fusion activities.

Here we report a distinct mechanism by which the Mfn2 catalytic function is regulated. Based on an unbiased mass spectrometry (MS) proteomic interactome analysis, we find that TAK1, a MAP3K family kinase (Shibuya et al., 1996), directly inhibits the GTP-binding capacity of Mfn2 through phosphorylation. Using molecular, cellular and *in vivo* approaches, we demonstrate that this new phosphoregulatory site has critical roles in mitochondrial remodeling and metabolism during development and tissue homeostasis.

## RESULTS

Our interest in understanding the complex TGF-β signaling and transcriptional networks prompted us to perform unbiased MS-based proteomic interactome screenings on all major TGF-β effectors including TAK1. As the focus of the present study, the TAK1 interactome analysis yielded Mfn2 as an intriguing and highly unexpected candidate (Figure S1). Their putative interaction was first validated by co-immunoprecipitation in both overexpression and endogenous systems (Figure 1A, B), followed by immunofluorescence to assess how TAK1 might affect the mitochondrial morphology. Here a brief treatment with various TAK1 activators including TGF-β, BMP9, TNF-α and LPS caused rapid mitochondrial fragmentation that increased over time in many epithelial, endothelial and fibroblast cell types (Figure 1C-E graphs; S2A). To confirm that these effects were TAK1-specific, we monitored for changes in mitochondrial morphology upon pharmacologic inhibition of the TAK1 kinase and found dramatically increased fusion under both basal and TGF-β-treated conditions (Figure 1F; graph). Similarly, silencing endogenous TAK1 expression abrogated the fission-inducing effects of TGF-β (Figure 1G graph; S2B), thus supporting a role for TAK1 in promoting mitochondrial fission through a physical interaction with Mfn2.

**Figure 1.**
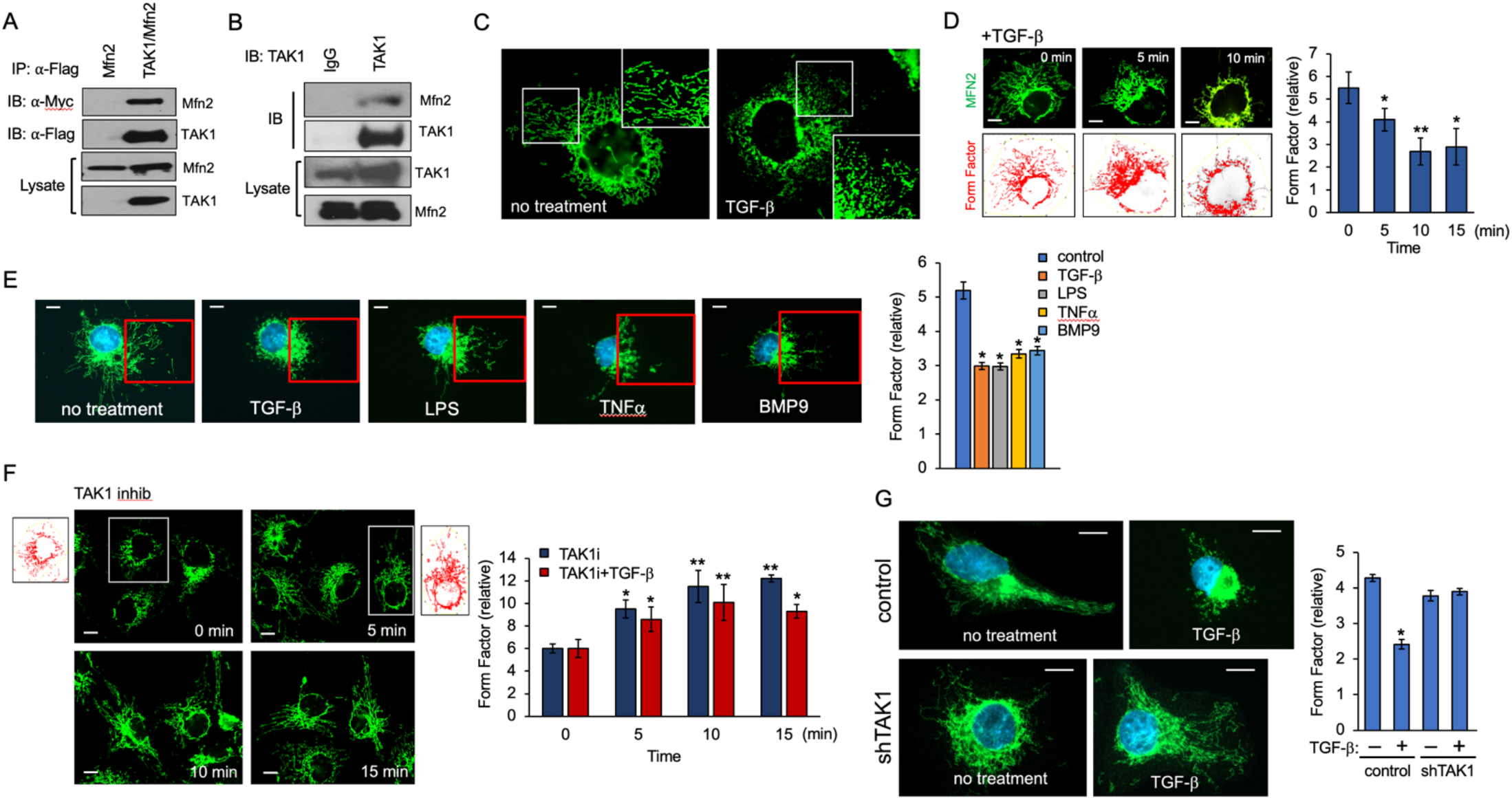
TAK1-Mfn2 interaction enhances mitochondrial fragmentation. (A) COS-7 cells expressing Myc-Mfn2 and Flag-TAK1 were immunoprecipitated with Flag antibody. (B) COS-7 cell lysate was immunoprecipitated with either IgG control or TAK1 antibody. (C) Representative immunofluorescence images of COS-7 cells serum starved for 2-3 h prior to treatment with TGF-β (200 pM) for 15 min, then stained for Mfn2 (green). (D) Immunofluorescence analysis of Mfn2 (upper panels in green) and Form Factor outlines (lower panels in red) in serum starved COS-7 cells treated with TGF-β (200 pM) for the indicated time points. Form Factor graph reflects the degree of mitochondrial fusion. At least 30 cells were quantified from 3 independent experiments. Error bars represent SEM. *p<0.05, **p<0.01 relative to control. (E) COS-7 cells serum starved for 2-3 h prior to treatment with TAK1 activators including TGFβ (200 pM), LPS (20 ng/mL), TNFα (20 ng/mL) and BMP-9 (50 ng/mL) for 30 min. Results are representative of 3 independent experiments with 20-30 images per group/experiment. Graph shows Form Factor values. *p<0.03 relative to control. (F) COS-7 cells cultured in growth media were treated with TAK1 inhibitor (5Z-7-Oxozeaneol 20 μM) for the indicated times in the presence or absence of TGFβ (200 pM). Graph represents Form Factor quantification is based on at least 30 cells per group. *p<0.05, **p<0.05 relative to control. (G) PANC1 WT and PANC1 shTAK1 cells were serum starved for 3 hours prior to treatment with TGFβ (200 pM) for 30 min. Images are representative of at least 3 independent experiments, and 30 images were chosen for form factor quantification. *p<0.05 relative to control.

A series of biochemical experiments were performed to test how TGF-β influences the TAK1-Mfn2 interaction and determine whether it is direct. First, consistent with TGF-β causing a real-time enhancement of mitochondrial fission, the TAK1-Mfn2 interaction increased when cells were briefly stimulated with TGF-β prior to their co-immunoprecipitation, as was also the case when co-expressed with TAB1, a TAK1 adaptor protein known to promote TAK1 activation (Figure 2A, B). Second, a protein binding assay using recombinant purified proteins derived from the baculovirus/insect cell system further confirmed that the Mfn2-TAK1 interaction was direct (Figure 2C). To identify the structural determinants of their interaction, we made deletions of the conserved Mfn2-GTPase domain (Mfn2^ΔGTPase^) and the C-terminal region (Mfn2^ΔCT^) (Figure 2D schematic). Here, TAK1 immunoprecipitation resulted in a similar level of Mfn2^ΔCT^ coprecipitation as WT, whereas Mfn2^ΔGTPase^ interaction was severely diminished, indicating that TAK1 binds to the Mfn2-GTPase domain (Figure 2D). A reciprocal TAK1 mutagenesis study revealed that the entire TAK1 C-terminal segment (TAK1^ΔCT^) was dispensable for Mfn2 binding (Figure 2E), thus demonstrating that the TAK1 kinase core likely engages the Mfn2-GTPase domain in a TGF-β-responsive manner.

**Figure 2.**
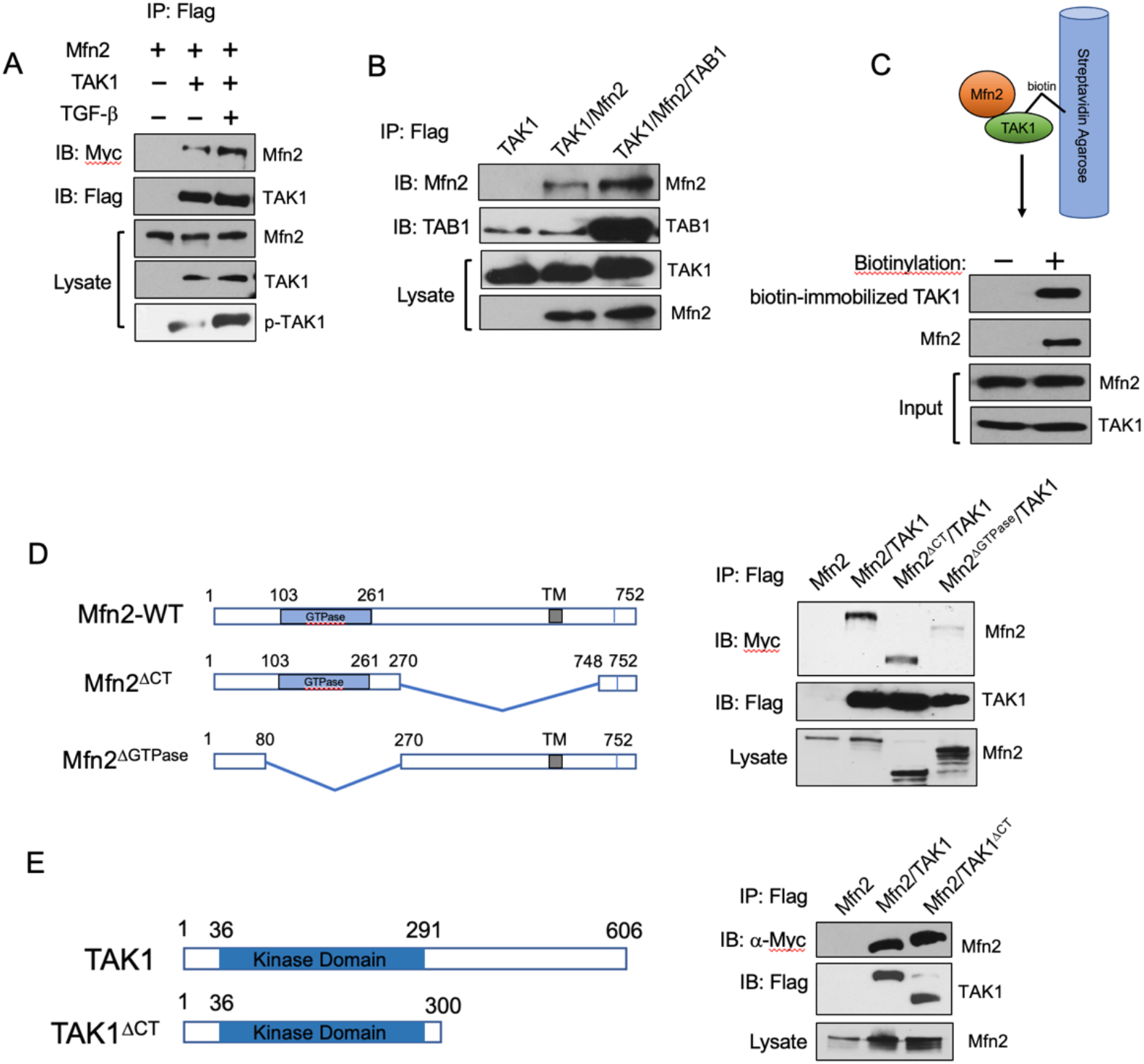
Structural requirements of the TAK1-Mfn2 interaction. (A) Western blot shows COS-7 cells expressing Flag-TAK1 and Myc-tagged Mfn2 were treated with TGF-β (200pM) for 30 min in the presence or absence of 5Z-7-Oxozeaneol (20μM) followed by Flag immunoprecipitation. (B) Western blot shows COS-7 cells expressing Flag-TAK1, TAB1 and/or Mfn2 were immunoprecipitated using Flag antibody. (C) Purified Mfn2 was incubated with purified and biotinylated TAK1 (0.5 μM each) immobilized to streptavidin agarose. As negative control, unlabeled Mfn2 and TAK1 were co-incubated in streptavidin agarose. (D) Schematic of Mfn2 WT and deletion mutants Δ^GTPase^ and Δ^CT^. COS-7 cells expressing Myc-Mfn2 WT, Δ^GTPase^, Δ^CT^ and Flag-TAK1 were immunoprecipitated using a Flag antibody. (E) Schematic of wildtype TAK1 and Δ^CT^. COS-7 cells expressing Myc-Mfn2 WT and Flag-TAK1 WT and Δ^CT^ were immunoprecipitated using a Flag antibody.

Based on above results, we hypothesized that TAK1 inhibits Mfn2 function through phosphorylation. To test this, we first compared the levels of mitochondrial fusion in cells overexpressing the wildtype, TAK1^ΔCT^ or TAK1^K63W^, a TAK1 kinase-dead point mutant as previously described (Kishimoto et al., 2000; Shah et al., 2018). Here results showed strong mitochondrial fragmentation in cells expressing the wildtype and TAK1^ΔCT^ but not TAK1^K63W^ compared to control (Figure 3A; graph). Accordingly, we tested whether TAK1 directly phosphorylates the Mfn2-GTPase domain via phosphoproteomic mapping. Specifically, we focused on Mfn2 modifications associated with wildtype TAK1 but not TAK1^K63W^ and found Ser249 as a distinct phosphorylation site (Figure S3). Because Ser249 is a conserved yet functionally uncharacterized site (Figure 3B), we sought to determine its role through ectopic expression of a phosphomimetic (S249E or D) or an alanine mutant (S249A) in Mfn2-null MEFs. Reconstitution of Mfn2 revealed that, relative to WT, S249A dramatically increased fusion, whereas both S249D and S249E yielded constitutive mitochondrial fragmentation (Figure 3C). Similar outcomes were observed in parallel studies where Mfn2 and TAK1 co-overexpression resulted in significant loss of fusion in Mfn2 WT but not S249A, which strongly resisted the fission-causing effects mediated by TAK1 phosphorylation (Figure 3D; graph). However, Mfn2-S249 phosphomimetics exhibited constitutive mitochondrial fragmentation whether or not TAK1 was overexpressed (Figure 3D; graph). Importantly, ectopic expression of the S249D mutant produced a dominant negative effect in COS7 cells as the overall percentage of cells with severely fragment mitochondria increased dramatically compared to the expression of the wildtype and S249A (Figure S4). These results combined provided strong evidence that TAK1 directly phosphorylates Mfn2-Ser249 to inhibit mitochondrial fusion.

**Figure 3.**
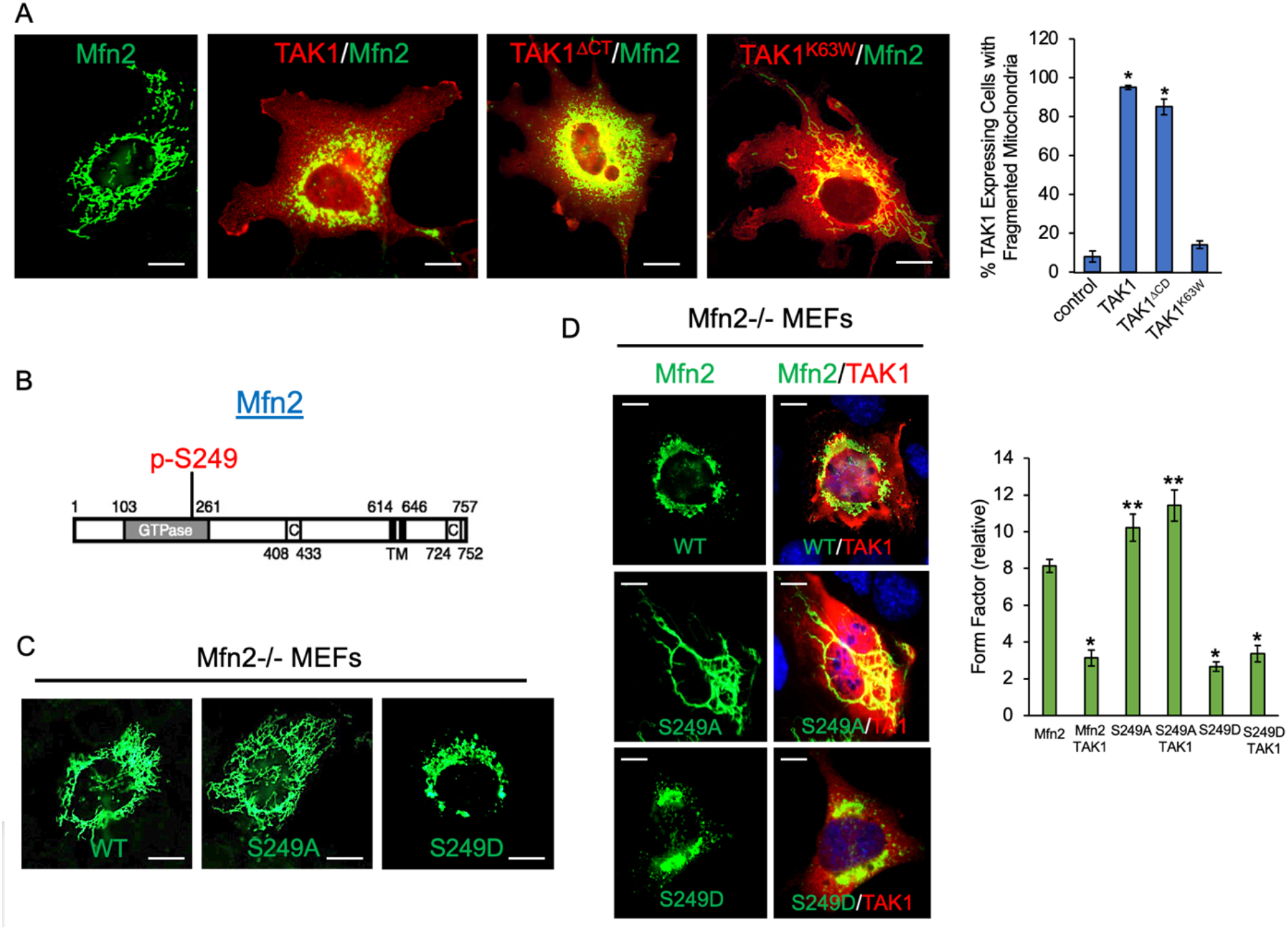
Identification of TAK1 phosphorylation site on Mfn2 GTPase domain. (A) MEF cells expressing Flag-TAK1, Flag-TAK1^ΔCD^ or Flag-TAK1^K63W^ were stained for Flag (red) and endogenous Mfn2 (green). Graph indicates percentage of Flag staining-positive cells (>30 cells per group) with fragmented mitochondrial morphology. Quantification and representative images are based on at least 3 independent experiments. *p<0.02 relative to control. (B) Schematic of the conserved S249 phosphorylation site (red) residing in the Mfn2 GTPase domain. (C) Mfn2-null MEF cells were ectopically rescued with Mfn2 WT, Mfn2-S249A or Mfn2-S249E prior to immunofluorescence staining for Mfn2 (green). (D) Representative immunofluorescence images of Mfn2-null MEFs ectopically expressing Myc-Mfn2 WT, S249A or S249E (green) with TAK1 (red). Graph quantification of Form Factor was based on analysis of at least 30 cells per group from 3 independent experiments. *p<0.02, **p<0.05 relative to Mfn2 alone.

To determine the molecular basis for how S249 phosphorylation inhibits fusion, we explored the possibility that this site modulates the GTP-binding properties of Mfn2 given that it resides within the GTPase domain. To test this hypothesis, we turned to a fluorescence-based Mfn2-GTP binding assay using MANT-GTP, a GTP analog which we previously showed enhances fluorescence emission upon selectively binding to a GTPase (Kumar et al., 2016). First, using purified proteins, we observed significant basal fluorescence enhancement in S249A compared to WT and S249D (Figure 4A). Similar binding patterns were observed when using immunoprecipitated Mfn2, where S249A displayed the highest MANT-binding while the S249D, much like the Mfn2^DGTPase^ mutant that is unable to bind GTP and thus used as negative control, was nearly devoid of fluorescence enhancement (Figure 4B). In replicate experiments using the immunoprecipitation approach, Mfn2 was co-expressed with various TAK1 mutants prior to Mfn2-immunoprecipitation. Here, Mfn2 displayed a marked increase in MANT-binding when isolated from cells overexpressing TAK1^K63W^, whereas those immunoprecipitated from WT and TAK1^ΔCT^ expressing cells had reduced binding compared to Mfn2 alone (Figure 4C). This TAK1 kinase-dependent inhibition of MANT-binding was further evident in Mfn2 WT upon TGF-β treatment but not when co-incubated with TAK1 inhibitor, whereas S249A and S249D immunoprecipitates exhibited little to no change in MANT-binding (Figure 4D), thus demonstrating that Mfn2-S249 acts as a key phosphoregulatory site of Mfn2 activation.

**Figure 4.**
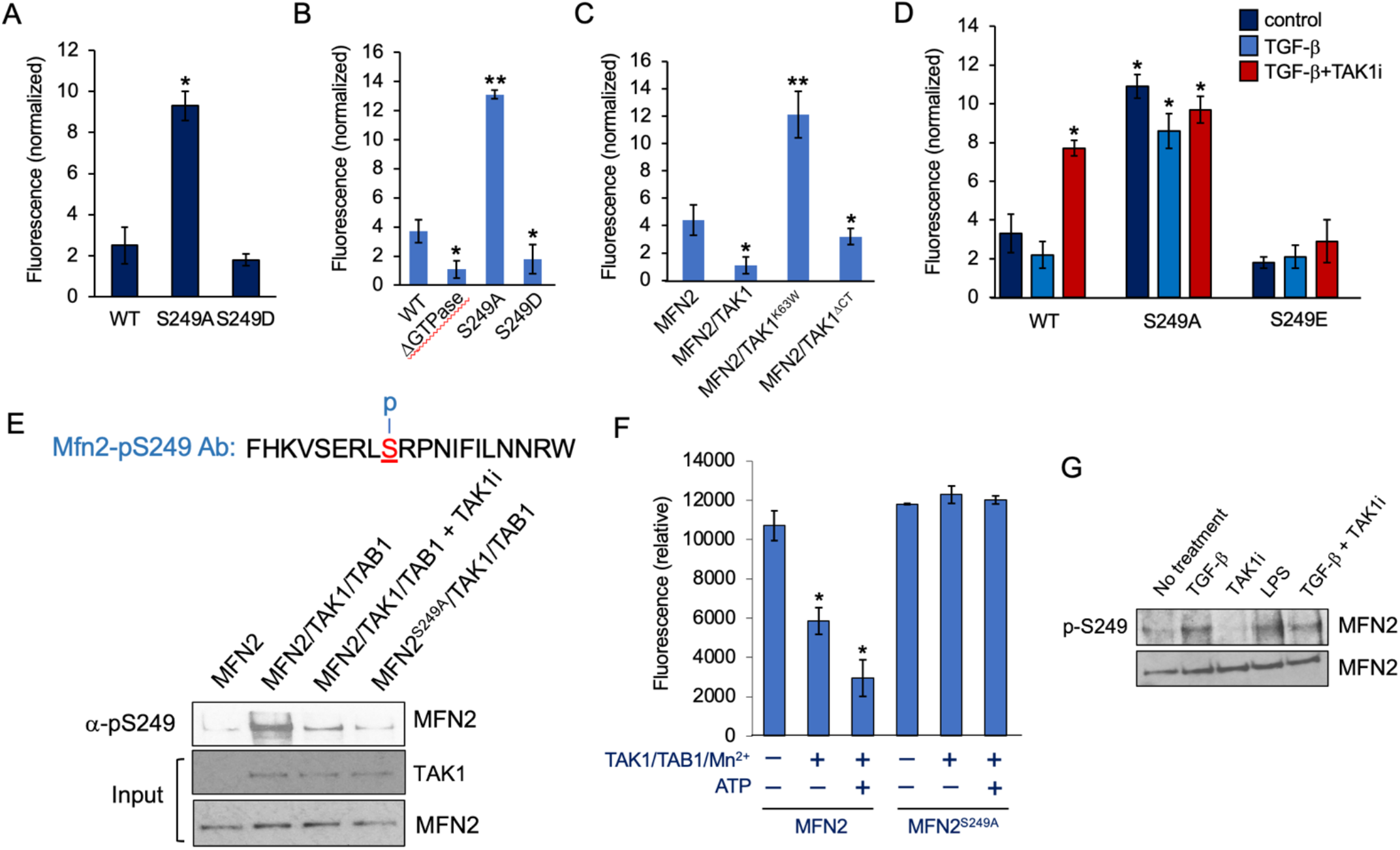
Mfn2-S249 phosphorylation by TAK1 inhibits GTP-binding. (A) Graph represents changes in MANT-GTP fluorescence emission upon binding purified Mfn2 WT, S249A or S249E (0.5 μM). Control MANT-GTP (2.5 μM) emission was subtracted from each condition, then normalized to Mfn2 alone plus MANT-GTP (2.5 μM). Error bars represent SE from 3 independent experiments. *p<0.001 relative to WT. (B) Graph represents MANT-GTP emission (2.5 μM) upon incubation with Mfn2 WT and indicated mutants ectopically expressed, then immunoprecipitated from Mfn2-null MEFs. Error bars represent SE from at least 3 independent experiments *p<0.01; **p<0.001 relative to WT. (C) Graph represents MANT-GTP emission of immunoprecipitated Mfn2 upon coexpression with TAK1, TAK1^K64W^ or TAK1 ^ΔCD^. Error bars represent SE from 3 independent experiments *p<0.03; **p<0.001 relative to Mfn2 alone. (D) Graph represents MANT-GTP emission upon incubation with immunoprecipitated Mfn2 WT, S249A or S249E from cells treated with TGF-β (200pM) in the presence or absence of 5Z-7-Oxozeaneol (20μM) for 30 min. Error bars represent SE from 3 independent experiments. *p<0.001 relative to WT no treatment. (E) Schematic (upper) represents the peptide sequence used to generate a phospho-specific Mfn2-S249 antibody. Western blot (lower) represents an *in vitro* kinase reaction using purified Mfn2 WT or Mfn2^S249^A (0.5 μM each) upon incubation with purified TAK1/TAB1 complex (0.5 μM) and Mg^2+^/ATP for 30 min in the presence or absence of 5Z-7-Oxozeaneol. (F) Graph represents MANT-GTP emission (2.5 μM) upon binding purified Mfn2 WT or MFN2^S429A^ that was incubated with purified TAK1/TAB1 and Mn^2+^ in the presence or absence of ATP (30 min). Error bars represent SE from at least 3 independent experiments *p<0.005 relative to Mfn2 WT alone. (G) Western blot shows p-Mfn2-S249 levels from whole cell lysates of PANC1 cells serum starved for 3 h prior to treatment with TGF-β (200pM), LPS (10 ng/mL) for 30 min or pretreated with 5Z-7-Oxozeaneol (20μM) for 15 min followed by TGF-β stimulation.

To provide direct evidence that TAK1 phosphorylates Mfn2 at Ser249 to inhibit GTP-binding, we generated a custom Ser249 phospho-specific antibody to detect Mfn2-S249 phosphorylation in an *in vitro* kinase reaction. Here, basal S249 phosphorylation was nearly undetectable in the purified Mfn2 when alone, while co-incubation with the TAK1/TAB1 complex caused a sharp increase in the p-S249 signal (Figure 4E). As expected, both pharmacologic TAK1 inhibition or the use of Mfn2-S249A blocked S249 phosphorylation, hence indicating that Mfn2 is a direct phosphosubstrate for TAK1 (Figure 4E). Moreover, to establish Mfn2-S249 as a key phosphoregulatory site targeted by TAK1 to modulate GTP-binding, a replicate *in vitro* kinase reaction was performed followed by immediate MANT-binding measurements. Here results showed strong S249 phosphorylation-induced loss of GTP-binding in the wildtype whereas Mfn2-A displayed constitutively high levels of MANT-fluorescence irrespective of TAK1 kinase activity (Figure 4F). While these *in vitro* findings explained how TAK1 can regulate Mfn2 activation at the molecular level, we further tested whether Mfn2-S249 phosphorylation occurs in response to TAK1 activation in intact cells. Indeed, in various cell types, both TGF-β and LPS treatment resulted in rapid increase in Mfn2-S249 phosphorylation whereas co-treatment with a TAK1 inhibitor abrogated this effect (Figure 4G).

We next examined how TAK1-dependent Mfn2 inactivation influences bioenergetics by measuring for changes in ATP and mitochondrial ROS production. In both normal and cancer cell types, a brief 30 min TGF-β stimulation caused a moderate reduction in ATP production relative to control, whereas TAK1 inhibition enhanced ATP production independent of TGF-β (Figure 5A). In contrast, the presence of mitochondrial ROS increased upon TGF-β treatment while TAK1 inhibition reduced the output (Figure 5B), thus indicating that TAK1-induced Mfn2 inactivation reduces bioenergetics while promoting oxidative stress.

**Figure 5.**
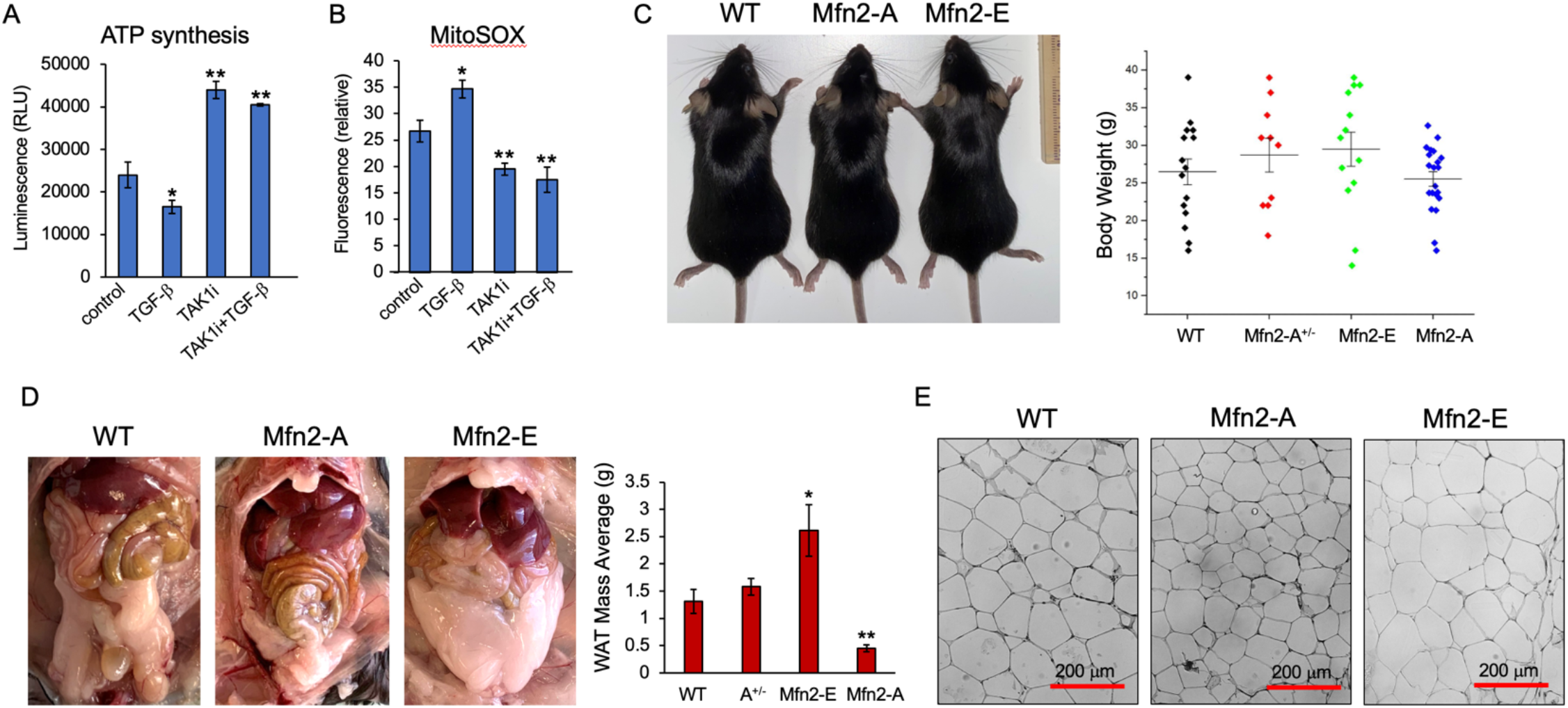
Cellular and *in vivo* effects of TAK1-induced Mfn2-S249 phosphorylation. (A) Graph represents luminescence as a measure of ATP production in COS-7 cells upon treatment with TGF-β1 (200 pM) for 30 min in the presence or absence of 5Z-7-oxozeaenol (20 μM). Error bars represent SEM. *p<0.05; **p<0.01 relative to control. (B) Graph represents mitochondrial superoxide levels in COS-7 cells upon treatment TGF-β (200 pM) for 30 min or pretreatment in the presence or absence of 5Z-7-oxozeaenol (20 μM). Error bars represent SEM and type 2 T-test shows relative to control: *p<0.04; **p<0.02 relative to control. (C) Representative images of 16 to 20 week-old male WT, S249A homozygote (Mfn2-A) and S249E heterozygous (Mfn2-E) mice. Graph show body weight (g) of WT, S249A heterozygote (Mfn2-A+/-), Mfn2-E and Mfn2-A. Error bars represent SEM. *p>0.05; no statistical difference among the groups compared to WT. (D) Representative images of visceral WAT in WT, Mfn2-A and Mfn2-E mice. Graph represents mass (g) of isolated adipose tissue from the mice. Error bars represent SEM: *p<0.04; **p<0.02 relative to WT. (E) Representative images of H&E staining of WAT sections derived from the indicated mice.

To gain further biological insights, we generated Mfn2-S249 knock-in point mutant mice using CRISPR-based genome-editing. Unexpectedly, we determined that S249E homozygous mutation causes embryonic lethality since repeated attempts even with various breeding strategies still failed to yield any litters, whereas S249A homozygotes (hereafter referred to as Mfn2-A) proved viable with no overt phenotypes other than being slightly smaller and leaner in overall body weight than WT (Figure 5C; graph). However, Mfn2-E heterozygotes (hereafter referred to as Mfn2-E) were viable but exhibited increased visceral fat accumulation upon weaning despite their similar body weight, food intake with standard diet (Figure 5C, D; graphs). Consistent with S249 phosphorylation in promoting the mass of white adipose tissues (WAT), Mfn2-A mice displayed the least amount of WAT, typically ranging from two to five-fold less fat than WT or Mfn2-E mice (Figure 5C, D graphs).

Upon ruling out caloric intake as a major underlying factor, we anticipated that behavioral or neuromuscular defects could contribute to the changes in adiposity. To address this, we first measured for plasma leptin levels as well as anxiety and cognition measurements and observed no significant changes in these mutant mice (Figure S5). Similarly, no discernable changes were found in voluntary or provoked movements, nor discernable changes in thermal and tactile sensations among the mutant mice compared to control (Figure S6). Given these null effects, we then focused on potential cell-autonomous effects of Mfn2-S249 phosphorylation. Here, H&E staining indicated that adipocyte populations in the epididymal fat of Mfn2-A mice had a reduced cell area compared with WT and Mfn2-E (Figure 5E). Furthermore, mature primary white adipocytes isolated from the mice yielded similar results, with Mfn2-A adipocyte populations being uniformly smaller compared to WT and Mfn2-E (Figure 6A; graph).

**Figure 6.**
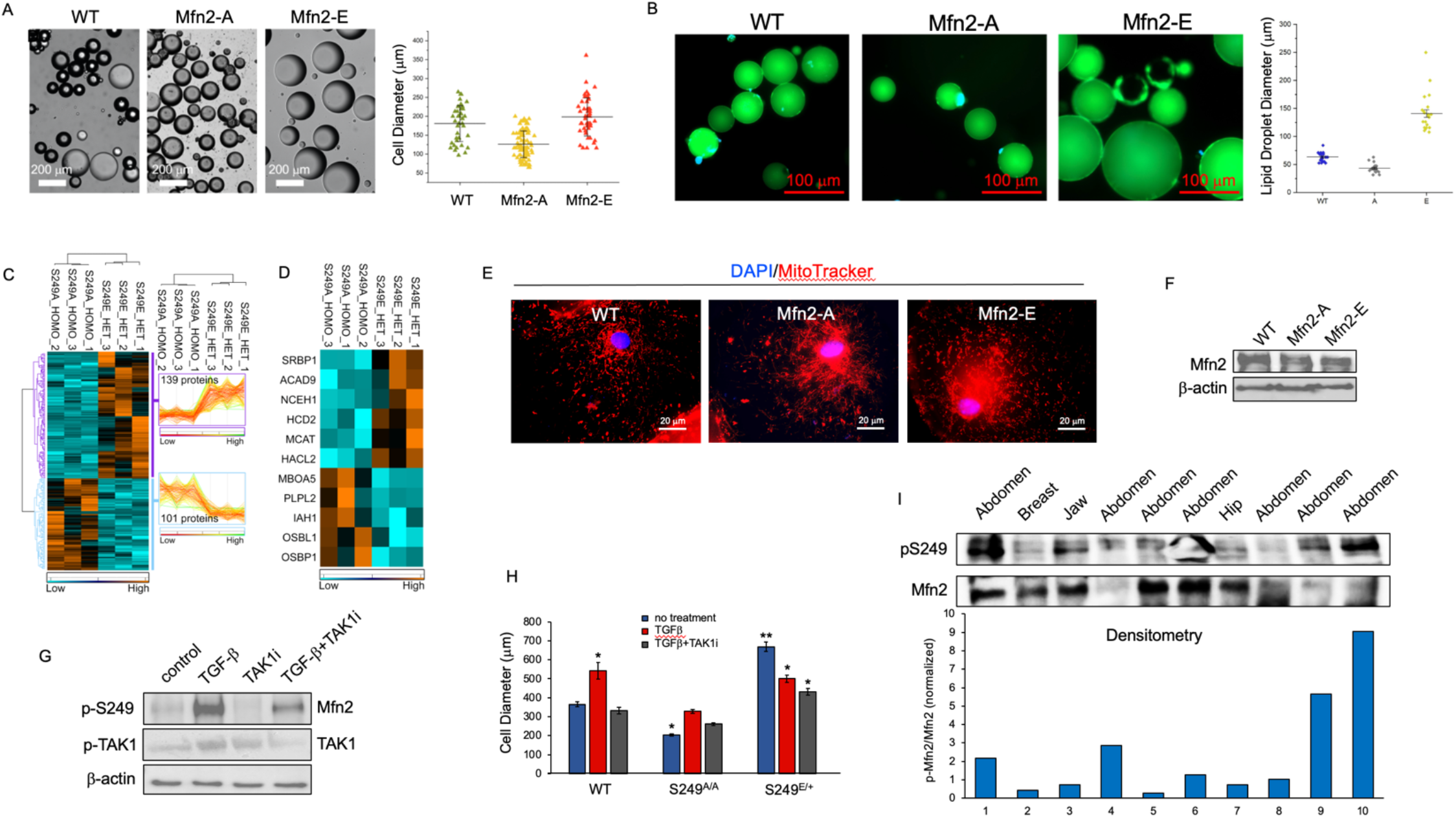
Mfn2-S249 phosphorylation-dependent changes in white adipocyte biology. (A) Representative images and graph quantification of the diameter of mature white adipocytes isolated from WT, Mfn2-A and Mfn2-E mice. Error bars represent SEM. p<0.05 for Mfn2-A and Mfn2-E groups relative to WT. (B) Representative images of BODIPY staining of lipid droplets (green) and nuclei (DAPI in blue) in primary adipocytes. Graph shows the diameter distribution of the droplets. p<0.05 for Mfn2-A and Mfn2-E groups relative to WT. (C) Quantitative proteomic analysis of WAT isolated from Mfn2-A versus Mfn2-E mice. A 2-way ANOVA was performed to identify significantly different protein levels between the two groups. Experiment was based on n=3 mouse WAT samples per group. (D) Quantitative proteomic data extracted from Figure 6C were subjected to further analysis based on lipid metabolism. (E) Immunofluorescence images represent primary white adipocytes stained with MitoTracker (red) and DAPI. (F) Western analysis shows Mfn2 protein levels in primary adipocytes isolated from the indicated mice. Cell lysates were normalized for protein level via Bradford Assay. (G) Western blot shows p-Mfn2-S249 levels in primary fat cells treated with TGF-β (200 pM) for 30 min in the presence or absence of 5Z-7-oxozeaenol (20 μM). (H) Graph represents quantification of the cell diameter of primary adipocytes isolated from mice upon treatment with TGF-β (200 pM) for 24 h in the presence or absence of 5Z-7-oxozeaenol (20 μM). Error bars represent SEM and p values relative to control: *p<0.04; **p<0.02. (I) Western blot shows p-Mfn2-S249 and total Mfn2 levels of cell lysates derived from various primary fat tissues isolated from patients who have undergone liposuction. Densitometry graph represents the relative p-Mfn2-S249 levels found in each patient sample.

As mature white adipocytes store fat as single lipid droplets that make up the majority of their cell size, we reasoned that the increased WAT mass in Mfn2-E is a result of adipocyte hypertrophy arising from enhanced lipid storage. As expected, BODIPY staining in primary adipocytes isolated from Mfn2-E mice showed significantly greater lipid droplet mass compared to that of Mfn2-A or WT (Figure 6B; graph). To explore the mechanisms underlying these changes in lipid mass, we performed a comparative unbiased quantitative proteomics analysis on primary mature adipocytes isolated from either Mfn2-A or Mfn2-E mice. Of the 5899 quantifiable proteins detected, 240 proteins were discovered to be significantly different between the two mutant strains, with 139 proteins exhibiting an increase in expression in Mfn2-E, while 101 proteins were decreased (Figure 6C). Of these significantly affected proteins, 11 were associated with lipid metabolism, many of which help explain the size difference between Mfn2-E and Mfn2-A adipocytes (Figure 6D). In particular, we noted that prominent lipases such as ATGL (PLPL2), which catalyzes the initial step in triglyceride hydrolysis in adipocyte lipid droplets, was more than 2.5-fold higher in Mfn2-A compared to Mfn2-E adipocytes while fatty acid synthases such as MCAT was more than 2.5-fold higher in Mfn2-E adipocytes (Figure 6D). In effect, given that WT primary adipocytes are closer in size to Mfn2-E than Mfn2-A, the results here suggest that the Mfn2-S249 phosphorylation level plays an important role in WAT homeostasis and that targeted inhibition of this phosphorylation promotes lipid metabolism in adipocytes.

While our proteomics data reveal that there are likely other contributing factors associated with adiposity besides enzymes related to lipid metabolism (Figure S7), more importantly, we sought to establish a decisive molecular link between TAK1-induced Mfn2 inactivation that promotes mitochondrial fission and fat cell size. To do so, we stained primary mature adipocytes with MitoTracker and found prominent mitochondrial fragmentation in Mfn2-E cells versus WT while Mfn2-A cells displayed hyperfused morphology (Figure 6E). Notably, because a recent study established the association between diminished fusion and Mfn2 protein levels with increasing fat deposition in WAT (Mancini et al., 2019), we then tested whether the Mfn2 expression is altered in the Mfn2 mutant mouse adipocytes. Consistent with when ectopically expressed, a direct comparison of Mfn2 expression in primary cells showed similar levels at basal state (Figure 6F), strongly suggesting that the observed changes in adipocyte size is linked to Mfn2-mediated fusion activity rather than total expression. To also test whether TAK1 inhibits Mfn2 activation in primary fat cells as observed in other cell types, we assessed for Mfn2-S249 phosphorylation levels in primary mature adipocytes to find a robust increase upon brief TGF-β stimulation, whereas TAK1 inhibition blocked this effect (Figure 6G).

To test whether TAK1 intrinsically promotes adipocyte hypertrophy, primary fat cells from WT and mutant mice were subjected to TGF-β treatment in the presence or absence of TAK1 inhibition (Figure 6H). Here we observed a profound increase in cell diameter of WT adipocytes upon TGF-β treatment over the course of 16 to 24 h whereas TAK1 inhibition using two different small molecule inhibitors abrogated this effect (Figure 6H; S8). In contrast, TGF-β failed to produce a strong response in lipid accumulation in Mfn2-A adipocytes irrespective of TAK1 inhibition, suggesting that the TGF-β-induced hypertrophy occurs primarily through TAK1 regulation of the S249 phosphoregulatory site (Figure 6H). However, there was an unexpected modest reduction in cell diameter of Mfn2-E adipocytes following TGF-β treatment over basal conditions (Figure 6H). Because the phosphomimetic substitution renders Mfn2-S249 phosphorylation irrelevant, there may be other compensatory mechanisms at play in limiting the TGF-β-induced effects on adipocyte growth. Finally, we evaluated how Mfn2-S249 phosphorylation levels correlate to adiposity in humans by surveying fat samples derived from patients undergoing liposuction procedures. Despite the limited sampling size, Mfn2-S249 phosphorylation was generally higher in abdominal visceral fat compared to breast, cheek or hip, thus indicating that TAK1-specific regulation of Mfn2 activation may be more selective and prevalent in visceral WAT than other fat types (Figure 6I).

## DISCUSSION

Defects in TGF-β signaling are closely linked to mitochondrial dysfunction in many malignant and metabolic disorders. Yet there have been limited advances in our understanding of its differential and often dichotomous effects–ranging from enhancing or inhibiting respiration to mitochondrial biogenesis and destruction during apoptosis (Casalena et al., 2012; Liu and Desai, 2015). While such effects are widely believed to mediated by Smad-dependent gene regulation (Ramesh et al., 2008; Ramesh et al., 2009; Sayeed et al., 2010; Yoon et al., 2005), whether TGF-β effectors can influence mitochondrial dynamics in real time remains fundamentally elusive. Our results demonstrate that TAK1, a major TGF-β activated kinase, induces mitochondrial fragmentation through a previously unknown Mfn2 phosphorylation site which, in effect, acts as a molecular switch of GTP-binding to control self-oligomerization and membrane fusion.

The direct interaction between TAK1 and Mfn2 is of significance with broad physiological implications. First, this finding suggests a new rationale for TAK1-targeted therapies in mitochondrial disorders as it bypasses the involvement of many other TAK1 downstream effectors including the MAPKs (ERK, p38 and JNK), AMPK and the NFkB pathway (Mukhopadhyay and Lee, 2020). Second, it remains a strong possibility that mitochondrial fusion is subject to regulation by numerous exogenous and endogenous factors that activate TAK1. Indeed, while our study involved only a limited group of TGF-β family ligands and inflammatory cytokines, there is a growing number of TAK-inducing conditions including the mitogen pathways, Wnt and integrin signaling, ceramides and even extracellular stressors such as osmotic stress, UV irradiation and hypoxia (Mukhopadhyay and Lee, 2020). How these individual factors contribute to the Mfn2 catalytic activity relative to induction of mitophagy or apoptosis remain to be investigated.

The molecular basis for how S249 phosphorylation actually inhibits GTP-binding is unclear. While it is feasible that the negative charge associated with the phosphorylation creates an electrostatic repulsion at the active site, our structural analysis of the GTPase domain of Mfn1 and Mfn2 argues against this possibility since S249 resides on the opposite side of the protein encompassing the GTP-binding pocket. Instead, S249 may function as a docking regulatory site for potential guanine exchange factors (GEFs) or GTPase-activating proteins (GAPs) as we previously reported (Kumar *et al*., 2016). Regardless, the ability of TAK1 to recognize this site raises the question of whether Mfn1 is also a phospho-target since Ser249 is largely conserved in Mfn1 (Mfn2: ERL<u>**S^249^**</u>RP versus Mfn1: ERL<u>**S^228^**</u>KP). While Mfn1 was not found in our MS-based TAK1 interactome screening, subsequent co-immunoprecipitation experiments confirmed their association (data not shown). However, determining whether Mfn1-S228 is targeted by TAK1 is not trivial given that their co-immunoprecipitation may be the result of heterodimerization with Mfn2. Moreover, since TAK1 notoriously lacks a consensus phosphorylation recognition motif, a more thorough investigation will be necessary to determine the molecular basis of the TAK1-Mfn1 interaction and their biological consequences.

While the fact that Mfn2-A homozygote mice are viable indicates that TAK1-induced Mfn2 phosphorylation is not essential during development, there are still distinct phenotypes associated with these mice which suggest an important role in tissue homeostasis. For instance, aside from their aforementioned leaner and smaller size, the Mfn2-A homozygote mice were more susceptible to developing spontaneous skin lesions and exhibited poor wound healing ability that may be due to defective immune responses and angiogenesis. Consistent with this notion, we observed intriguing differences in cytokine production in macrophages versus T-cells isolated from the mutant mice. Similarly, the abnormal patterning of the adult retinal vasculature observed in the mutant mice compared to WT further implicates a role for TAK1 regulation of Mfn2 activity in vascular remodeling (data not shown). In contrast, the embryonic lethality observed in Mfn2-E homozygotic mice indicates that persistent inhibition of GTP-binding may be just as deleterious as Mfn2-deficiency. While beyond the scope of this study, it will be important to examine how Mfn2-S249 phosphorylation is involved in maintaining homeostasis across different tissue types during aging and under various pathologic conditions.

It is also noteworthy that Mfn2-S249 phosphorylation has such a prominent effect on WAT. One potential explanation is that changes in mitochondrial morphology upon Mfn2 dysfunction are amplified in WAT than other cell types due to their lower abundance of Mfn1 and Mfn2 expression (Mancini *et al*., 2019). Along these lines, it is interesting that the effects of TGF-β on the size of lipid droplets in primary adipocytes are almost entirely dependent on TAK1 activity rather than the previously reported Smad3-mediated gene regulation of lipogenic enzymes since TAK1 inhibition abrogated the growth-promoting effects (Lin et al., 2009; Yadav et al., 2011). Results here suggest that, at least in mature white adipocytes, TAK1 may have a more significant role than Smads in driving the hypertrophic behavior. Moreover, this notion is further consistent with the higher Mfn2-S249 phosphorylation levels detected in visceral fat versus that of subcutaneous fat in human samples. While the overall sampling size was rather limited due to the difficulty of acquiring sufficient number of patient samples, we have demonstrated that the phospho-Mfn2-S249 antibody can serve as a potential biomarker of TAK1-specific mitochondrial defects in obesity and other human diseases.

Based on these results, we conclude that TAK1 is a key regulator of mitochondrial dynamics. It inhibits fusion through phosphorylation-induced Mfn2 catalytic inactivation, a process which we propose is crucial for metabolic homeostasis across many cell types independent of TGF-β/Smad-induced mitochondrial biogenesis and cell death programs.

## ACKNOWLEDGMENTS

We thank the University of Arizona Cancer Center for their assistance. This work was supported in part by the internal funding provided by the University of Arizona Cancer Center and College of Medicine.

## AUTHOR CONTRIBUTIONS

S.K. conducted the majority of the experiments and data analysis as first author; A.R., T.A. and E.K., P.C.F., N.S., H.R.O. performed biochemical experiments and generated stable cell lines. K.L.K., T.W.V. and T.M.L.M performed mouse behavioral studies. T.G.G. generated the CRISPR knock-in mice. P.R.L. provided MS proteomics data analysis. D.T.H. provided the patient fat samples. Y.S.L. and K.M. provided assistance with experimental procedures and edited the paper. N.Y.L. conceived the study and edited the paper with contributions from S.K.

---

**Figure S1.**
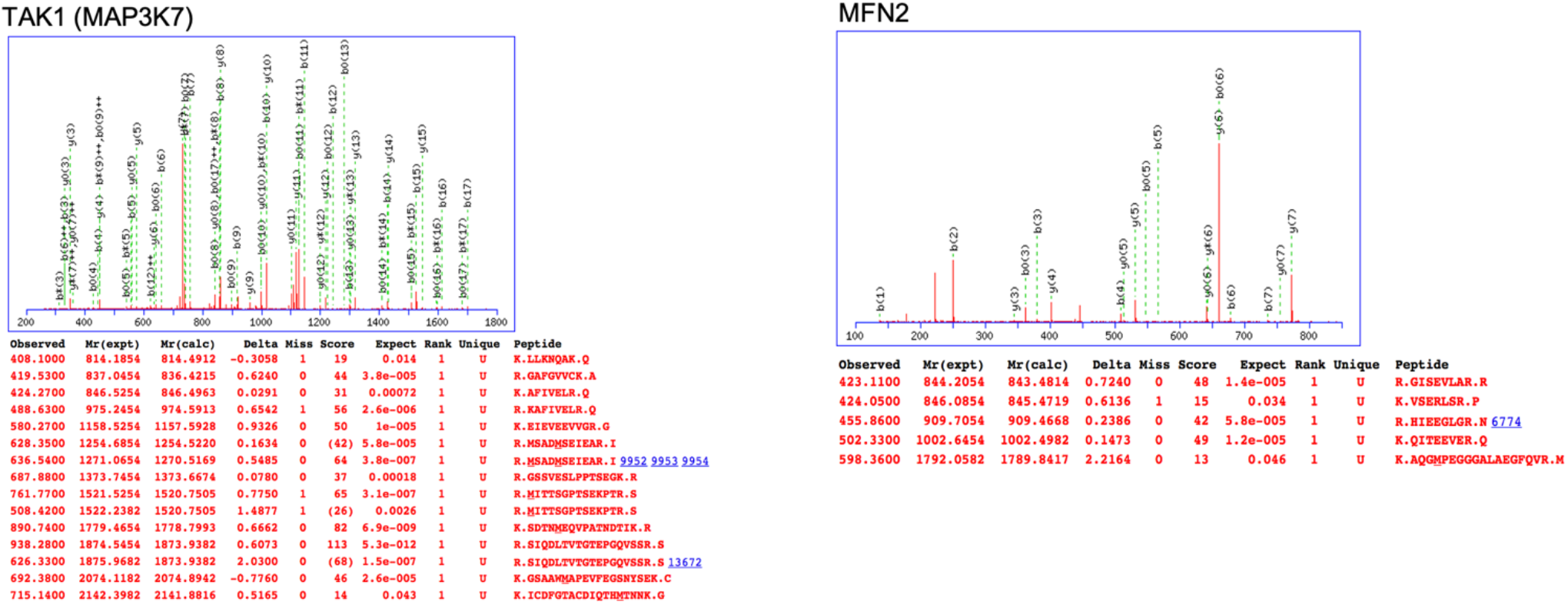
Identification of Mfn2 as a novel binding partner for TAK1 through MS proteomics analysis. Shown are the MS peptide fragment identification of TAK1 and Mfn2. Immunoprecipitation of Flag-TAK1 in COS-7 cells were subjected to nano LC-MS/MS and proteomics analysis. The mass spectrometry chromatogram peaks and internal sequence of the protein that matched to peptide fragments for TAK1 and Mfn2 are shown in red.

**Figure S2.**
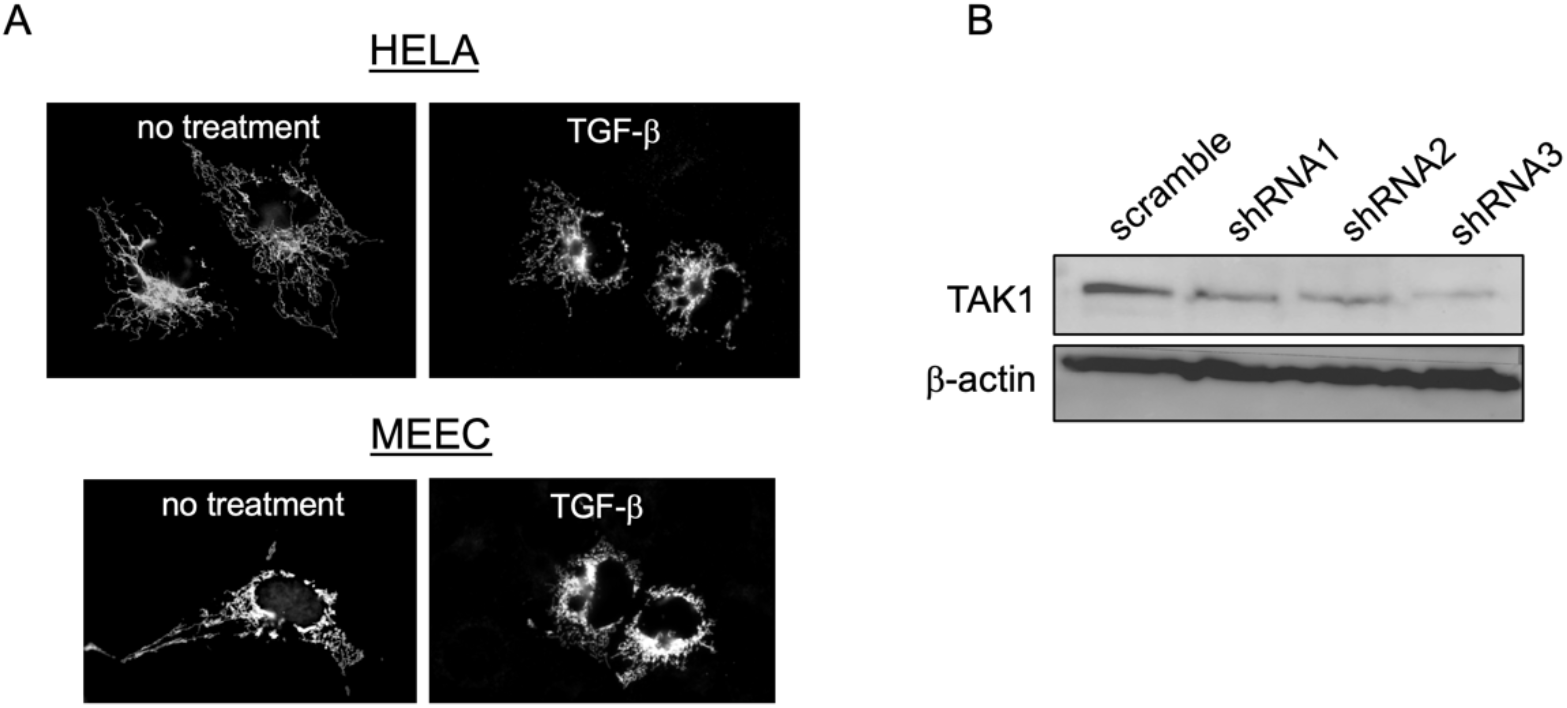
Effects of TGF-β-induced TAK1 activation on mitochondria morphology. (A) Shown are immunofluorescence images of HeLA and mouse embryonic endothelial cells (MEECs) serum starved for 3 h prior to TGF-β stimulation (200 pM) for 30 min. (B) Western blot shows endogenous TAK1 depletion using three different shRNA against human TAK1 in PANC1 cells. **Figure S3**. MS-phosphoproteomics spectral analysis

**Figure S3.**
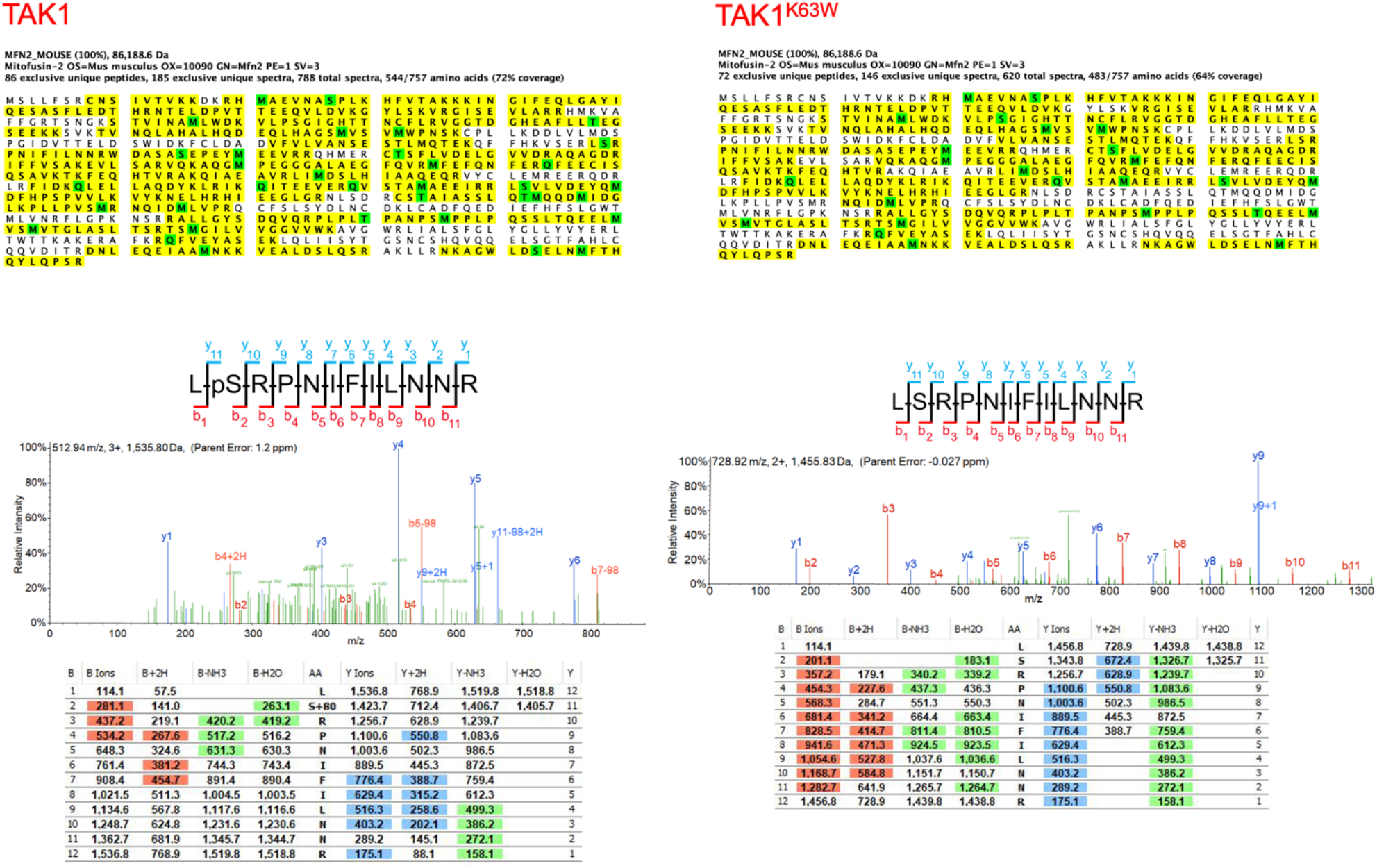
TAK1-dependent Mfn2-S249 phosphorylation. Myc-Mfn2 was transiently transfected with either TAK1 WT or TAK1^K63W^ in COS-7 cells. Mfn2 was then subjected to nano LC-MS/MS and proteomics analyses upon immunoprecipitation. Shown are representative spectral peaks indicating phosphorylation signals of S249 when Mfn2 was co-expressed with TAK1 WT but not TAK1^K63W^.

**Figure S4.**
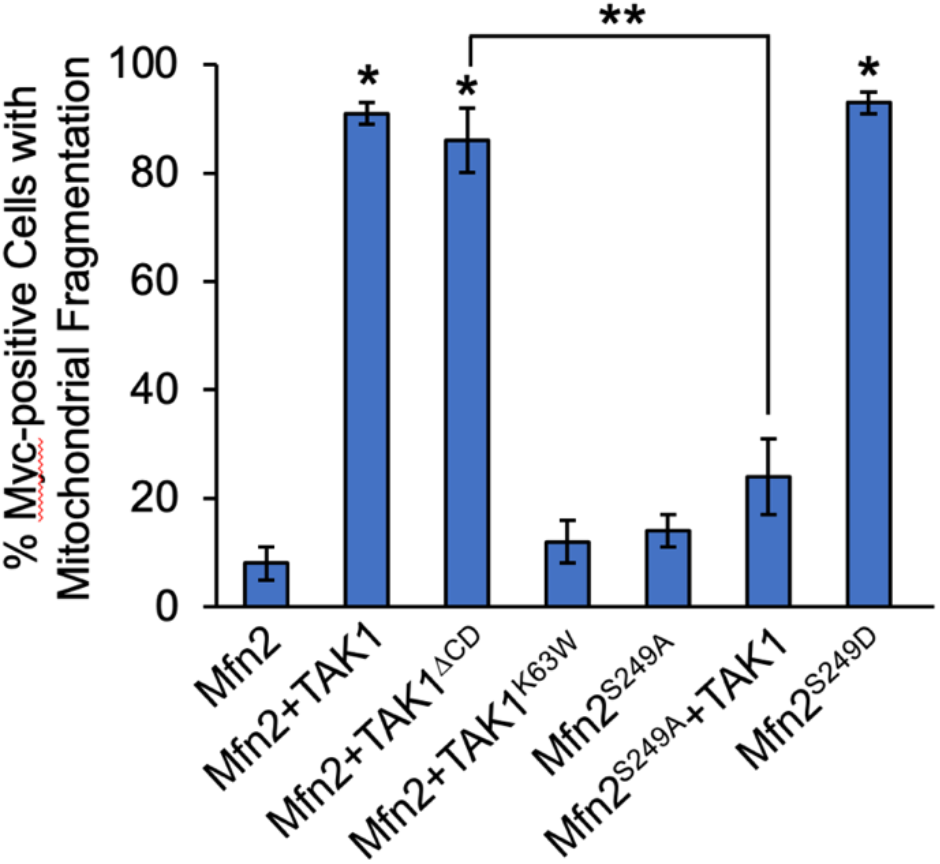
Effects of TAK1-dependent Mfn2-S249 phosphorylation on mitochondrial fusion. MFN2-null MEFs were transiently transfected with either Myc-Mfn2, TAK1 WT, TAK1^ΔCD^, TAK1^K63W^, Mc-Mfn2^S249A^ or Myc-Mfn2^S249D^ prior to fixation and immunofluorescence staining for Myc (green) and Flag (red). Graph quantification shows percentage of Myc-positive cells with fragmented mitochondrial morphology. At least 30 cells were scored per group from at least 3 independent experiments. p<0.01 relative to Mfn2 alone; **p<0.02 as indicated.

**Figure S5.**
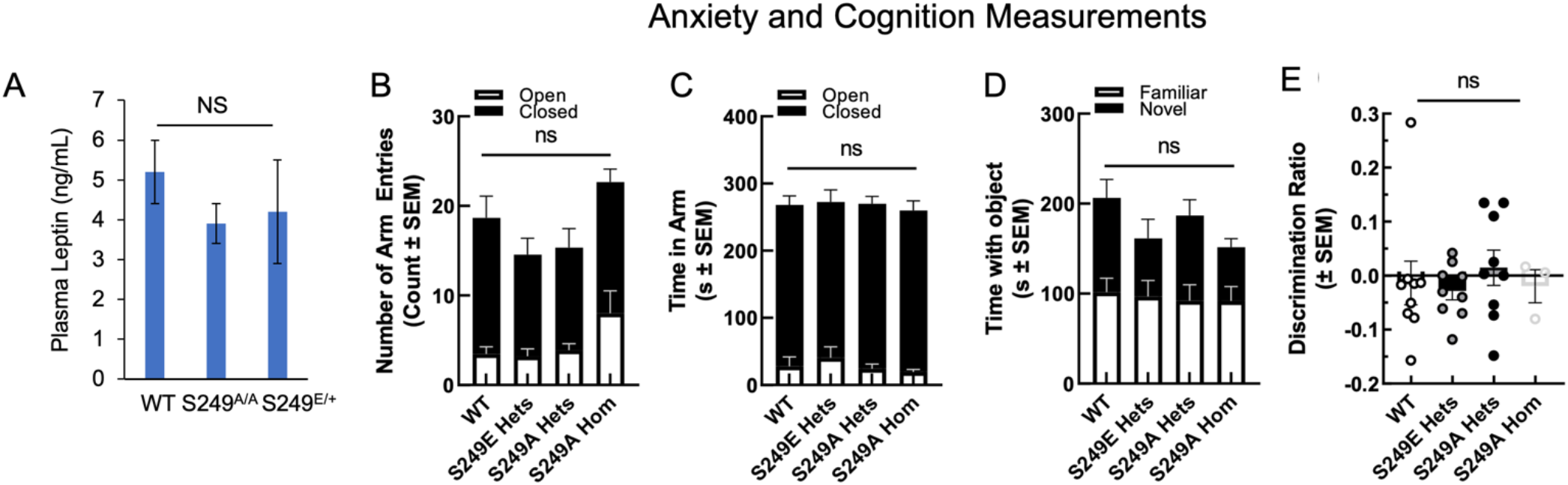
Leptin, anxiety and cognition Measurements. **(A)** Serum leptin levels were measured using Quantikine Leptin ELISA assay. N=3 mice per group. Mfn2-S249 mutation did not significantly alter (B) number of closed arm entries or (C) time spent in each arm during testing on the elevated plus maze for anxiety. Cognitive outcomes of time spent with object (D) and the discrimination ratio (E) during evaluation of novel object recognition were not changed.

**Figure S6.**
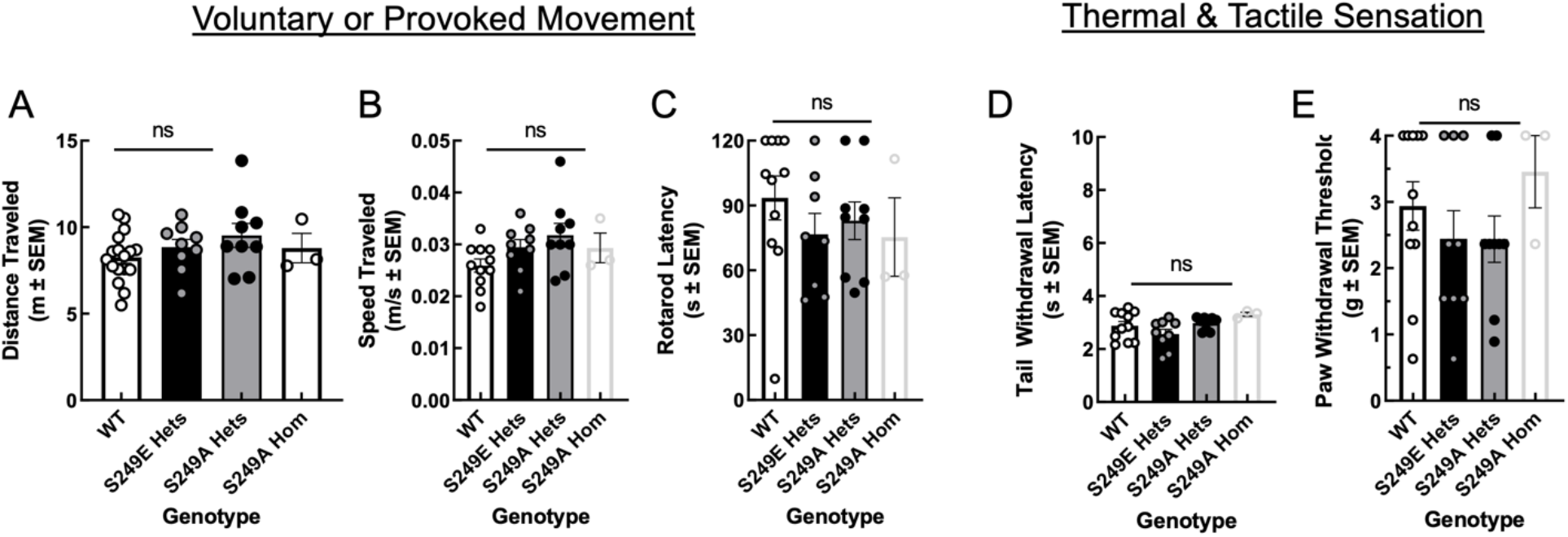
Movement and thermal/tactile sensation measurements. Mfn2-S249 mutation did not reduce the ability to move as determined by distance traveled (A) and speed (B) during exploration in open field or provoked movement on a Rotarod (C). Thermal (D) and tactile (E) sensations were also intact.

**Figure S7.** Quantitative proteomics Raw Excel file.

**Figures S8.**
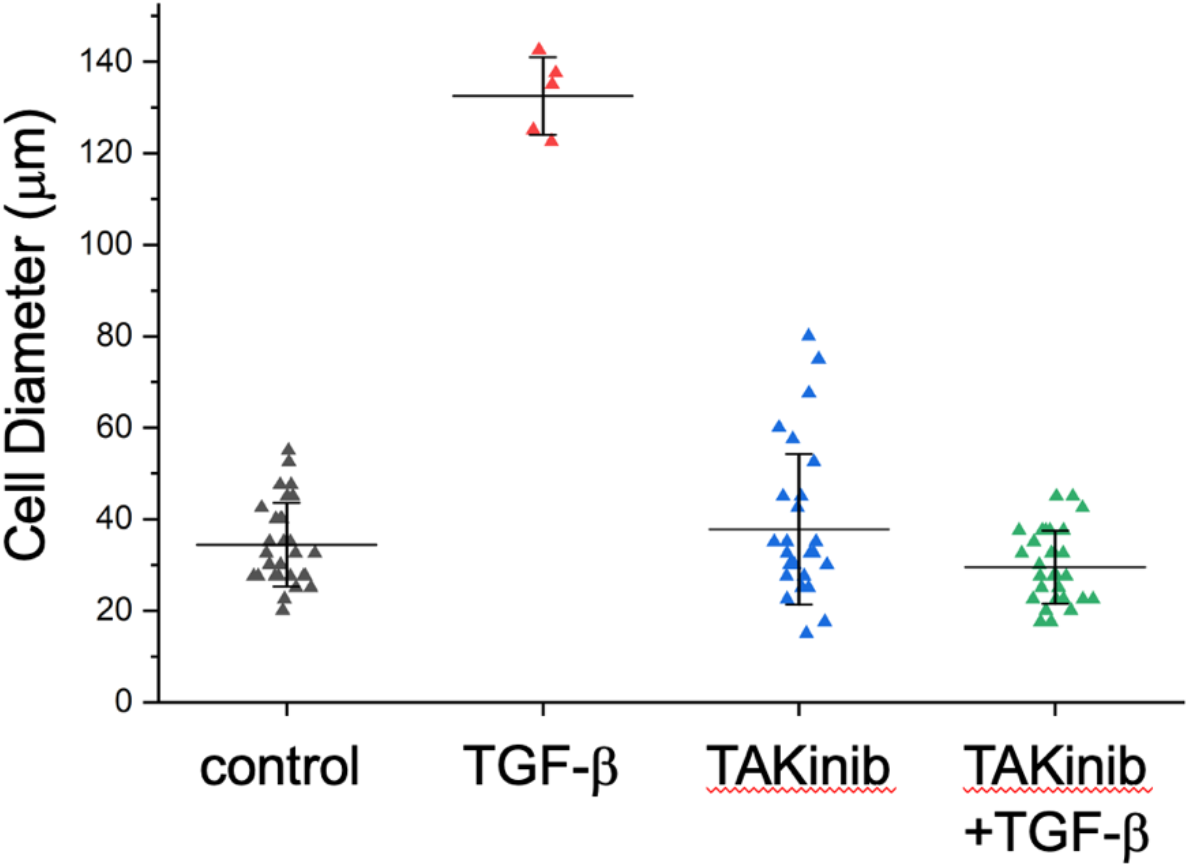
TAK1 inhibition abrogates TGF-β-induced increase in lipid storage. Graph shows quantification of the size (diameter) of primary fat cells upon treatment with TGF-β (200 pM) for 30 min or TAKinib (20 uM) for 15 min or pretreatment with TAKinib (20uM) for 15 min followed by TGF-β (200 pM) stimulation for 30 min. Error bars represent SEM.

## Method Details

### Cell Culture

COS-7, PANC-1 WT and PANC-1 shTAK1 were cultured in DMEM (GIBCO) with 10% fetal bovine serum (FBS) supplement (GIBCO). PANC-1 shTAK1 stables were selected using puromycin (10 ug/mL) after small hairpin RNA vector transfection using Lipofectamine 2000 (Invitrogen). Cells were grown in 5% CO_2_ and at 37°C.

### Immunoprecipitation

Cell lines were put on ice, washed with 1XPBS and treated with lysis buffer for 20 minutes (20 mM HEPES [pH 7.4], 150 mM NaCl, 2 mM EDTA, 10 mM NaF, 10% [w/v] glycerol, 1% NP-40). The cells were collected in a microcentrifuge tube and centrifuged at 13,000 RPM for 15 minutes. The supernatants were collected, and the appropriate antibodies were added prior to incubation. Agarose beads protein G or protein A were added for 4-6 hours at 4°C. The beads containing the immunoprecipitants were washed three times and stored in a mixture with 2X sample buffer for western blot analyses.

### Generation of recombinant baculoviruses and purified proteins

Genes encoding full-length human TAK1 and Mfn2 were fused with a 5’ hexahistidine-tag by PCR and then cloned into pFastBac1 vector (Life Technologies). Once constructed, each plasmid was transformed into DH10Bac competent cells for production of recombinant bacmids as we previously described (Kumar *et al*., 2016; Lee et al., 2006). Purified recombinant viruses were amplified and harvested. Sf21 cells infected with these viruses at 2 MOI were harvested 48 h later, then subjected to protein purification (Lee *et al*., 2006).

### Isolation of Primary White Adipocytes from Mice

White adipocytes were collected from WT, Mfn2-A and Mfn2-E mice. Adipocyte tissues were finely chopped and then transferred to collagenase solution (1.5mg/mL collagenase and 0.5% BSA in 1x DMEM media filtered with 0.2 μm syringe filter) for enzymatic digestion for 1 hour at 37°C. The digested fat tissue solution was filtered through a 300 μm cell strainer. The strained and washed tissue solution was centrifuged at 800 RPM for 1 min. The white fat layer floating on the top of the solution was collected and was transferred to a new tube with growth media containing 1x DMEM (4.5 g/L glucose), 10% FBS, 1% Pen streptomycin and was centrifuged again for two more times. After the last centrifugation the top fat layer was collected and plated on gelatin coated tissue culture plates for further biochemical analysis.

### Protein Extraction from Primary White Adipocytes

White adipocytes were collected from 27 weeks old male WT, Mfn2-A and Mfn2-E mice. Fat tissues were finely chopped and then homogenized in RIPA buffer (Tris HCl (pH 7.4) 20mM, EDTA 1Mm, EGTA 0.5Mm, Triton-X 1%, Na-deoxycholate 0.1%, SDS 0.1% and NaCl 150mM; Protease and Phosphatase Inhibitor 1:1000) by sonicating the tissue samples three times for 5 minutes on ice until the tissues were completely homogenized. The homogenized fat tissues were incubated on ice for 1 hour. The tissue samples were then centrifuged at 4°C, 20000g for 15 minutes. The fat layer (top layer) was carefully removed and the remaining supernatant was centrifuged again for three to four times until the maximum amount of fat had been removed. The lysates were normalized for protein level via Bradford Assay and the normalized samples were loaded into the SDS PAGE gels for western blot analysis.

### ATP synthesis assay

Cellular ATP synthesis was measured using an ATP Determination Kit (Invitrogen). Briefly, cells in different treatment conditions were resuspended in a reaction buffer, mixed with the ATP standard solution in a 96-well plate, and measured for luciferase luminescence at 560 nm. The ATP content in the samples was calculated from the standard curve.

### Measurement of Mitochondrial ROS

Mitochondrial superoxide production was measured using the MitoSOX Red mitochondrial superoxide indicator fluorescent probe (Invitrogen). Cells grown and treated with TGFβ and TAK1 inhibitors in 96-well plates were washed twice with PBS and subsequently incubated for 10 min with Mitosox Red (5μM) at 37°C. After the incubation, fluorescence was measured with a microplate reader set to 510 nm excitation 580 nm emission wavelengths.

### Measurement of circulating leptin

Average plasma leptin concentrations were measured from each age-matched mouse (n=3) using the R&D Systems Quantikine ELISA Mouse/Rat Leptin Immunoassay according to manufacture instructions.

### Immunofluorescence

Cells grown on coverslips were fixed with 4% paraformaldehyde, permeabilized in 0.1% Triton X-100 in PBS for 3 min, then blocked with 5% BSA in PBS for 20 min. All primary and fluorescently conjugated secondary antibodies and DAPI were incubated at room temperature for 1 hr unless noted otherwise. Mitochondrial morphology assessment was based on calculating form factor, an average of isolated mitochondrial particles in a region of interest. Raw images obtained from immunofluorescence microscope were binarized and quantified based on ImageJ using Mito Morphology Macro (Dagda et al., 2009) and the equation Pm2/4π × Am, where Pm is the length of mitochondrial outline and Am is the area of mitochondrion (Mortiboys et al., 2008).

### Adipose Tissue Sample Preparation for H&E Staining

After extraction o f Adipose tissue, tissue was fixed in 4% paraformaldehyde overnight, the following day the tissue was washed 2 times with 1x PBS for 10 min, followed by storage in 70% ethanol at 4°C. the sample was stored for a couple days prior to being sent to IHC staining facility.

### Mass spectrometry and Database Search

HPLC-ESI-MS/MS was performed in positive ion mode on a Thermo Scientific Orbitrap Fusion Lumos tribrid mass spectrometer fitted with an EASY-Spray Source (Thermo Scientific, San Jose, CA). NanoLC was performed exactly as previously described (Kruse et al., 2017; Parker et al., 2019). Tandem mass spectra were extracted from Xcalibur ‘RAW’ files and charge states were assigned using the ProteoWizard 3.0 msConvert script using the default parameters. The fragment mass spectra were searched against the mus musculus SwissProt_2018_01 database (16965 entries) using Mascot (Matrix Science, London, UK; version 2.6.0) using the default probability cut-off score. The search variables that were used were: 10 ppm mass tolerance for precursor ion masses and 0.5 Da for product ion masses; digestion with trypsin; a maximum of two missed tryptic cleavages; variable modifications of oxidation of methionine and phosphorylation of serine, threonine, and tyrosine. Cross-correlation of Mascot search results with X! Tandem was accomplished with Scaffold (version Scaffold_4.8.7; Proteome Software, Portland, OR, USA). Probability assessment of peptide assignments and protein identifications were made using Scaffold. Only peptides with ≥ 95% probability were considered. Label-free peptide/protein quantification and identification. Progenesis QI for proteomics software (version 2.4, Nonlinear Dynamics Ltd., Newcastle uponTyne, UK) was used to perform ion-intensity based label-free quantification as previously described (Uhlorn et al., 2021).

### MitoTracker Staining of Primary Adipocytes

After isolation white adipocytes, a fraction of the white fat layer was transferred to a 5ml centrifuge tube and washed once with fresh growth media (1x DMEM (4.5 g/L glucose), 10% FBS, 1% Pen streptomycin) followed by incubation of 100mM mitotracker for 30min. Adipocytes were then washed with growth media before fixation with 4% PFA at RT for 20 min followed by 3x washes with 50mM glycine/ 1xPBS, aspirate the infranatant after centrifugation at 210g for 30s. Adipocytes were then incubated in 5μg/ml Hoechst for 10 min at RT for nuclear staining, cells were then plated on a cover slip in antifade solution for microscopy.

### Cognitive and Behavioral Studies

Male and female mice were assessed for motor skills, sensory thresholds, anxiety, and cognition as previously reported (Hay et al., 2019; Levine et al., 2020; Sandweiss et al., 2017; Vekariya et al., 2020).

## KEY RESOURCES TABLE

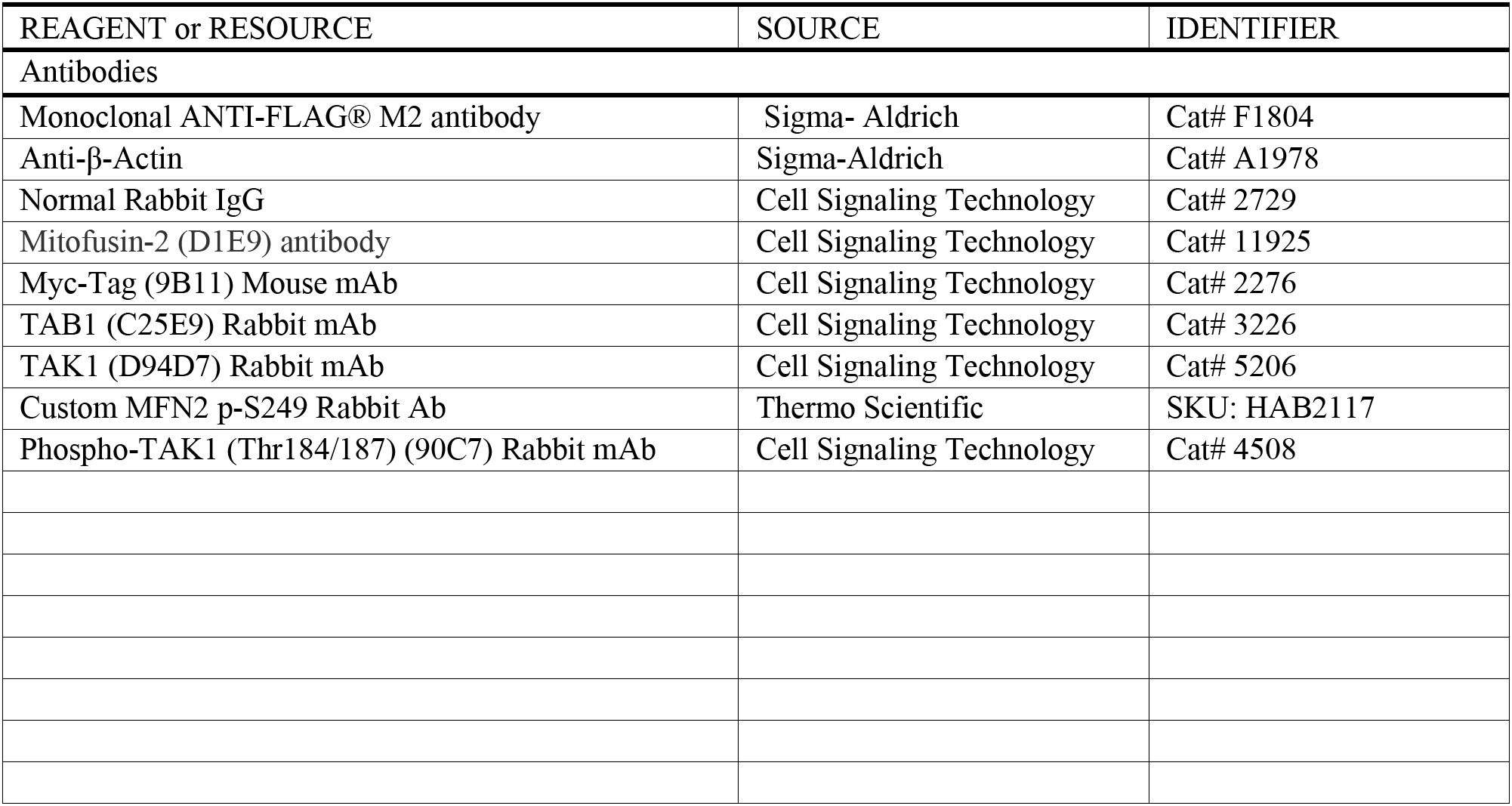
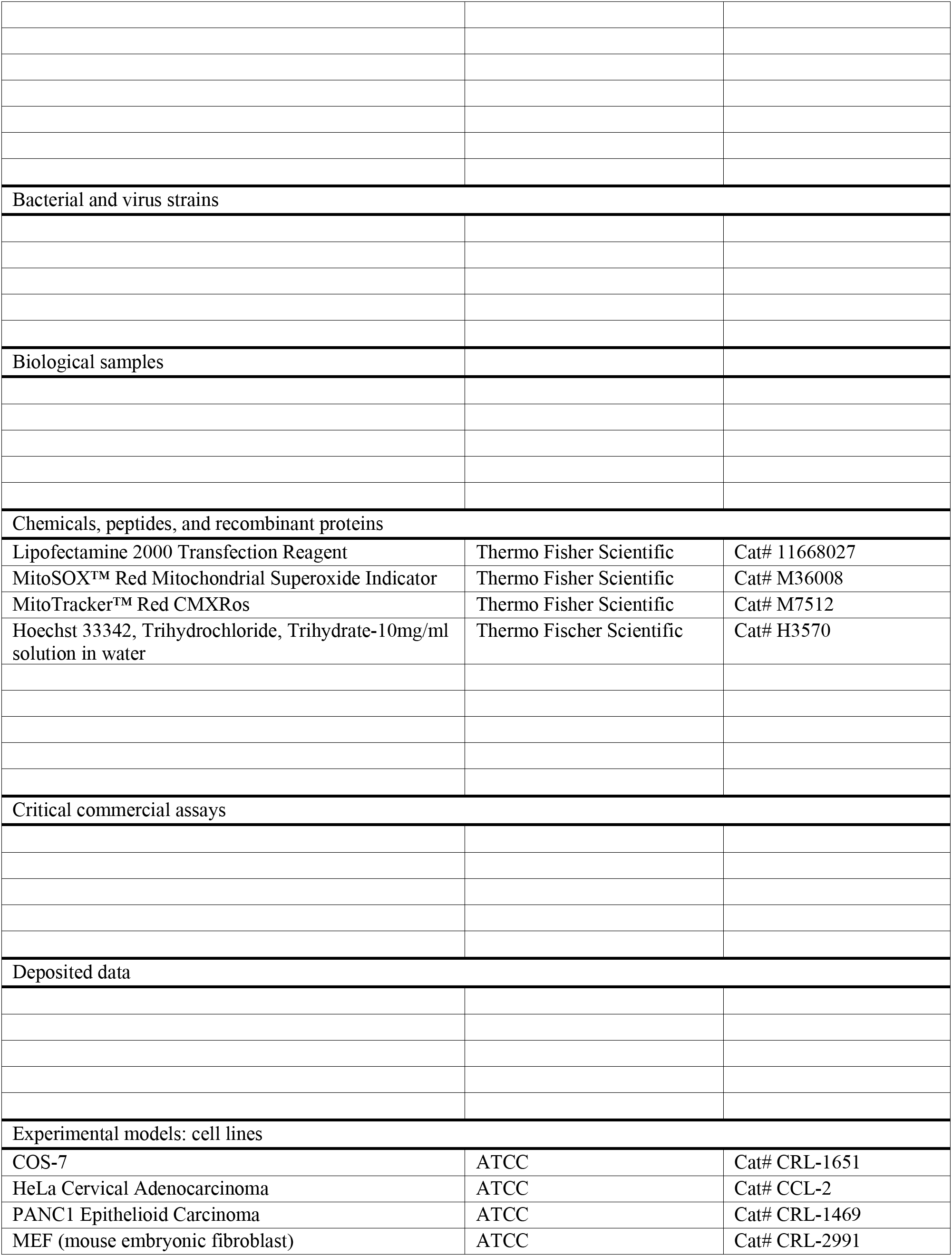
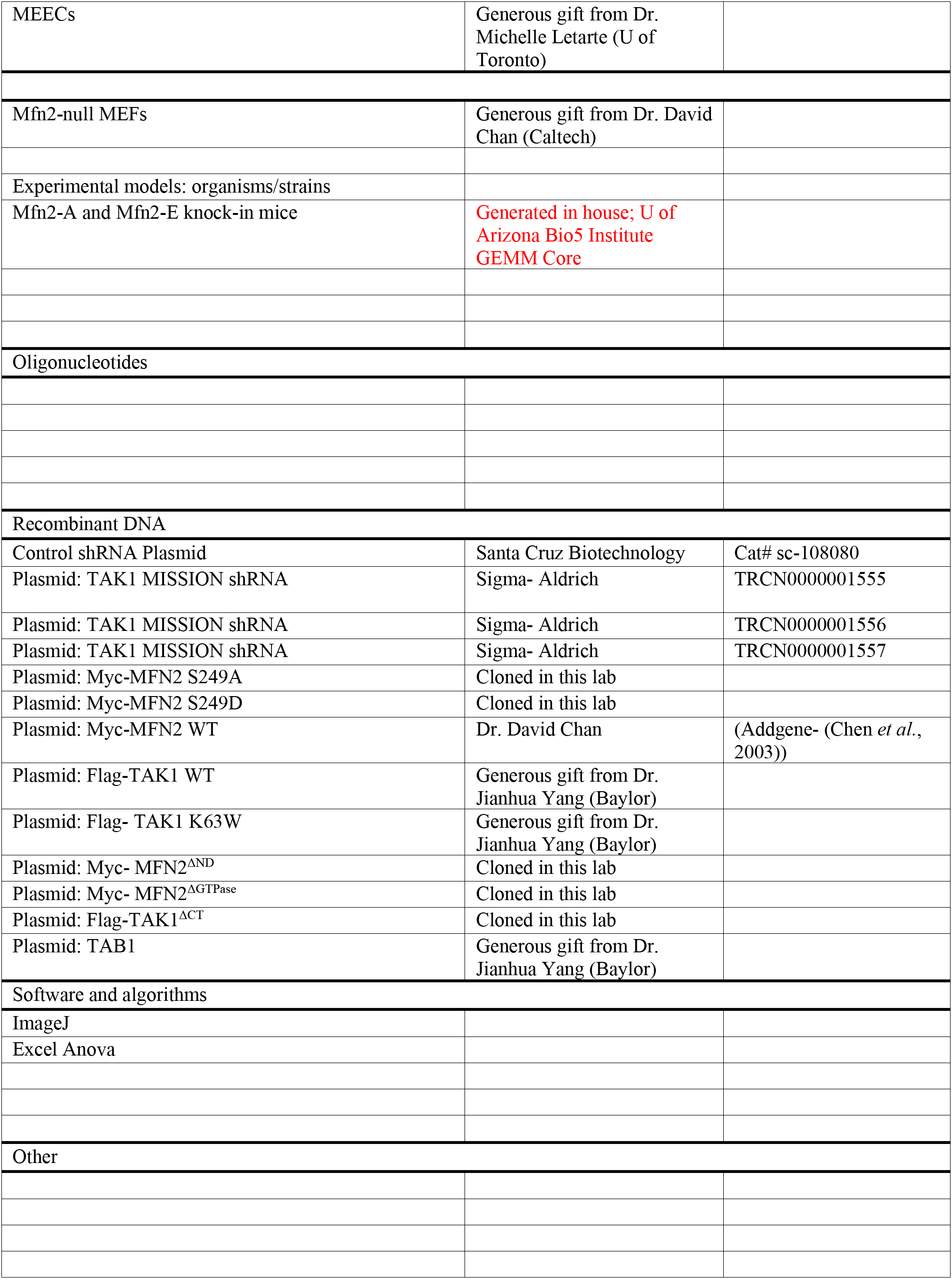

## REFERENCES

Cao, Y.L., Meng, S., Chen, Y., Feng, J.X., Gu, D.D., Yu, B., Li, Y.J., Yang, J.Y., Liao, S., Chan, D.C., and Gao, S. (2017). MFN1 structures reveal nucleotide-triggered dimerization critical for mitochondrial fusion. Nature 542, 372–376. 10.1038/nature21077.

Casalena, G., Daehn, I., and Bottinger, E. (2012). Transforming growth factor-beta, bioenergetics, and mitochondria in renal disease. Semin Nephrol 32, 295–303. 10.1016/j.semnephrol.2012.04.009.

Chen, H., Detmer, S.A., Ewald, A.J., Griffin, E.E., Fraser, S.E., and Chan, D.C. (2003). Mitofusins Mfn1 and Mfn2 coordinately regulate mitochondrial fusion and are essential for embryonic development. J Cell Biol 160, 189–200. 10.1083/jcb.200211046.

Chen, Y., and Dorn, G.W., 2nd (2013). PINK1-phosphorylated mitofusin 2 is a Parkin receptor for culling damaged mitochondria. Science 340, 471–475. 10.1126/science.1231031.

Dagda, R.K., Cherra, S.J., 3rd, Kulich, S.M., Tandon, A., Park, D., and Chu, C.T. (2009). Loss of PINK1 function promotes mitophagy through effects on oxidative stress and mitochondrial fission. J Biol Chem 284, 13843–13855. 10.1074/jbc.M808515200.

Eisner, V., Picard, M., and Hajnoczky, G. (2018). Mitochondrial dynamics in adaptive and maladaptive cellular stress responses. Nat Cell Biol 20, 755–765. 10.1038/s41556-018-0133-0.

Giacomello, M., Pyakurel, A., Glytsou, C., and Scorrano, L. (2020). The cell biology of mitochondrial membrane dynamics. Nat Rev Mol Cell Biol 21, 204–224. 10.1038/s41580-020-0210-7.

Hay, M., Polt, R., Heien, M.L., Vanderah, T.W., Largent-Milnes, T.M., Rodgers, K., Falk, T., Bartlett, M.J., Doyle, K.P., and Konhilas, J.P. (2019). A Novel Angiotensin-(1-7) Glycosylated Mas Receptor Agonist for Treating Vascular Cognitive Impairment and Inflammation-Related Memory Dysfunction. J Pharmacol Exp Ther 369, 9–25. 10.1124/jpet.118.254854.

Kishimoto, K., Matsumoto, K., and Ninomiya-Tsuji, J. (2000). TAK1 mitogen-activated protein kinase kinase kinase is activated by autophosphorylation within its activation loop. J Biol Chem 275, 7359–7364. 10.1074/jbc.275.10.7359.

Kruse, R., Krantz, J., Barker, N., Coletta, R.L., Rafikov, R., Luo, M., Hojlund, K., Mandarino, L.J., and Langlais, P.R. (2017). Characterization of the CLASP2 Protein Interaction Network Identifies SOGA1 as a Microtubule-Associated Protein. Mol Cell Proteomics 16, 1718–1735. 10.1074/mcp.RA117.000011.

Kumar, S., Pan, C.C., Shah, N., Wheeler, S.E., Hoyt, K.R., Hempel, N., Mythreye, K., and Lee, N.Y. (2016). Activation of Mitofusin2 by Smad2-RIN1 Complex during Mitochondrial Fusion. Mol Cell 62, 520–531. 10.1016/j.molcel.2016.04.010.

Leboucher, G.P., Tsai, Y.C., Yang, M., Shaw, K.C., Zhou, M., Veenstra, T.D., Glickman, M.H., and Weissman, A.M. (2012). Stress-induced phosphorylation and proteasomal degradation of mitofusin 2 facilitates mitochondrial fragmentation and apoptosis. Mol Cell 47, 547–557. 10.1016/j.molcel.2012.05.041.

Lee, N.Y., Hazlett, T.L., and Koland, J.G. (2006). Structure and dynamics of the epidermal growth factor receptor C-terminal phosphorylation domain. Protein Sci 15, 1142–1152. 10.1110/ps.052045306.

Levine, A., Liktor-Busa, E., Karlage, K.L., Giancotti, L., Salvemini, D., Vanderah, T.W., and Largent-Milnes, T.M. (2020). DAGLalpha Inhibition as a Non-invasive and Translational Model of Episodic Headache. Front Pharmacol 11, 615028. 10.3389/fphar.2020.615028.

Li, Y.J., Cao, Y.L., Feng, J.X., Qi, Y., Meng, S., Yang, J.F., Zhong, Y.T., Kang, S., Chen, X., Lan, L., et al. (2019). Structural insights of human mitofusin-2 into mitochondrial fusion and CMT2A onset. Nat Commun 10, 4914. 10.1038/s41467-019-12912-0.

Lin, H.M., Lee, J.H., Yadav, H., Kamaraju, A.K., Liu, E., Zhigang, D., Vieira, A., Kim, S.J., Collins, H., Matschinsky, F., et al. (2009). Transforming growth factor-beta/Smad3 signaling regulates insulin gene transcription and pancreatic islet beta-cell function. J Biol Chem 284, 12246–12257. 10.1074/jbc.M805379200.

Liu, R.M., and Desai, L.P. (2015). Reciprocal regulation of TGF-beta and reactive oxygen species: A perverse cycle for fibrosis. Redox Biol 6, 565–577. 10.1016/j.redox.2015.09.009.

Mancini, G., Pirruccio, K., Yang, X., Bluher, M., Rodeheffer, M., and Horvath, T.L. (2019). Mitofusin 2 in Mature Adipocytes Controls Adiposity and Body Weight. Cell Rep 27, 648. 10.1016/j.celrep.2019.03.065.

Mukhopadhyay, H., and Lee, N.Y. (2020). Multifaceted roles of TAK1 signaling in cancer. Oncogene 39, 1402–1413. 10.1038/s41388-019-1088-8.

Olichon, A., Baricault, L., Gas, N., Guillou, E., Valette, A., Belenguer, P., and Lenaers, G. (2003). Loss of OPA1 perturbates the mitochondrial inner membrane structure and integrity, leading to cytochrome c release and apoptosis. J Biol Chem 278, 7743–7746. 10.1074/jbc.C200677200.

Park, Y.Y., Nguyen, O.T., Kang, H., and Cho, H. (2014). MARCH5-mediated quality control on acetylated Mfn1 facilitates mitochondrial homeostasis and cell survival. Cell Death Dis 5, e1172. 10.1038/cddis.2014.142.

Parker, S.S., Krantz, J., Kwak, E.A., Barker, N.K., Deer, C.G., Lee, N.Y., Mouneimne, G., and Langlais, P.R. (2019). Insulin Induces Microtubule Stabilization and Regulates the Microtubule Plus-end Tracking Protein Network in Adipocytes. Mol Cell Proteomics 18, 1363–1381. 10.1074/mcp.RA119.001450.

Pyakurel, A., Savoia, C., Hess, D., and Scorrano, L. (2015). Extracellular regulated kinase phosphorylates mitofusin 1 to control mitochondrial morphology and apoptosis. Mol Cell 58, 244–254. 10.1016/j.molcel.2015.02.021.

Qi, Y., Yan, L., Yu, C., Guo, X., Zhou, X., Hu, X., Huang, X., Rao, Z., Lou, Z., and Hu, J. (2016). Structures of human mitofusin 1 provide insight into mitochondrial tethering. J Cell Biol 215, 621–629. 10.1083/jcb.201609019.

Ramesh, S., Qi, X.J., Wildey, G.M., Robinson, J., Molkentin, J., Letterio, J., and Howe, P.H. (2008). TGF beta-mediated BIM expression and apoptosis are regulated through SMAD3-dependent expression of the MAPK phosphatase MKP2. EMBO Rep 9, 990–997. 10.1038/embor.2008.158.

Ramesh, S., Wildey, G.M., and Howe, P.H. (2009). Transforming growth factor beta (TGFbeta)-induced apoptosis: the rise & fall of Bim. Cell Cycle 8, 11–17. 10.4161/cc.8.1.7291.

Sandweiss, A.J., Azim, A., Ibraheem, K., Largent-Milnes, T.M., Rhee, P., Vanderah, T.W., and Joseph, B. (2017). Remote ischemic conditioning preserves cognition and motor coordination in a mouse model of traumatic brain injury. J Trauma Acute Care Surg 83, 1074–1081. 10.1097/TA.0000000000001626.

Sayeed, A., Meng, Z., Luciani, G., Chen, L.C., Bennington, J.L., and Dairkee, S.H. (2010). Negative regulation of UCP2 by TGFbeta signaling characterizes low and intermediate-grade primary breast cancer. Cell Death Dis 1, e53. 10.1038/cddis.2010.30.

Shah, N., Kumar, S., Zaman, N., Pan, C.C., Bloodworth, J.C., Lei, W., Streicher, J.M., Hempel, N., Mythreye, K., and Lee, N.Y. (2018). TAK1 activation of alpha-TAT1 and microtubule hyperacetylation control AKT signaling and cell growth. Nat Commun 9, 1696. 10.1038/s41467-018-04121-y.

Shibuya, H., Yamaguchi, K., Shirakabe, K., Tonegawa, A., Gotoh, Y., Ueno, N., Irie, K., Nishida, E., and Matsumoto, K. (1996). TAB1: an activator of the TAK1 MAPKKK in TGF-beta signal transduction. Science 272, 1179–1182.

Smirnova, E., Griparic, L., Shurland, D.L., and van der Bliek, A.M. (2001). Dynamin-related protein Drp1 is required for mitochondrial division in mammalian cells. Mol Biol Cell 72, 2245–2256.

Uhlorn, J.A., Husband, N.A., Romero-Aleshire, M.J., Moffett, C., Lindsey, M.L., Langlais, P.R., and Brooks, H.L. (2021). CD4(+) T Cell-Specific Proteomic Pathways Identified in Progression of Hypertension Across Postmenopausal Transition. J Am Heart Assoc 10, e018038. 10.1161/JAHA.120.018038.

Vekariya, R.H., Lei, W., Ray, A., Saini, S.K., Zhang, S., Molnar, G., Barlow, D., Karlage, K.L., Bilsky, E.J., Houseknecht, K.L., et al. (2020). Synthesis and Structure-Activity Relationships of 5’-Aryl-14-alkoxypyridomorphinans: Identification of a mu Opioid Receptor Agonist/delta Opioid Receptor Antagonist Ligand with Systemic Antinociceptive Activity and Diminished Opioid Side Effects. J Med Chem 63, 7663–7694. 10.1021/acs.jmedchem.0c00503.

Yadav, H., Quijano, C., Kamaraju, A.K., Gavrilova, O., Malek, R., Chen, W., Zerfas, P., Zhigang, D., Wright, E.C., Stuelten, C., et al. (2011). Protection from obesity and diabetes by blockade of TGF-beta/Smad3 signaling. Cell Metab 14, 67–79. 10.1016/j.cmet.2011.04.013.

Yoon, Y.S., Lee, J.H., Hwang, S.C., Choi, K.S., and Yoon, G. (2005). TGF beta1 induces prolonged mitochondrial ROS generation through decreased complex IV activity with senescent arrest in Mv1Lu cells. Oncogene 24, 1895–1903. 10.1038/sj.onc.1208262.

Youle, R.J., and van der Bliek, A.M. (2012). Mitochondrial fission, fusion, and stress. Science 337, 1062–1065. 10.1126/science.1219855.

Zhou, W., Chen, K.H., Cao, W., Zeng, J., Liao, H., Zhao, L., and Guo, X. (2010). Mutation of the protein kinase A phosphorylation site influences the anti-proliferative activity of mitofusin 2. Atherosclerosis 211, 216–223. 10.1016/j.atherosclerosis.2010.02.012.

